# Shared components of heritability across genetically correlated traits

**DOI:** 10.1101/2021.11.25.470021

**Authors:** Jenna Lee Ballard, Luke Jen O’Connor

## Abstract

Most disease-associated genetic variants are pleiotropic, affecting multiple genetically correlated traits. Their pleiotropic associations can be mechanistically informative: if many variants have similar patterns of association, they may act via similar pleiotropic mechanisms, forming a shared component of heritability. We developed Pleiotropic Decomposition Regression (PDR) to identify shared components and their underlying genetic variants. We validated PDR on simulated data and identified limitations of existing methods in recovering the true components. We applied PDR to three clusters of 5-6 traits genetically correlated with coronary disease, asthma, and type II diabetes respectively, producing biologically interpretable components. For CAD, PDR identified components related to BMI, hypertension and cholesterol, and it clarified the relationship among these highly correlated risk factors. We assigned variants to components, calculated their posterior-mean effect sizes, and performed out-of-sample validation. Our posterior-mean effect sizes pool statistical power across traits and substantially boost the correlation (r^2^) between true and estimated effect sizes compared with the original summary statistics: by 94% and 70% for asthma and T2D out of sample, and by a predicted 300% for CAD.

## Introduction

Most disease-associated genetic variants affect multiple traits pleiotropically^1,2,3,4,5,6^, and these variants may act through pleiotropic mechanisms, shared across traits. Often, multiple pleiotropic variants – perhaps acting through similar pleiotropic mechanisms – have correlated effect sizes, leading to a nonzero genetic correlation^3,5^. Occasionally, these correlations arise when one trait is causal for the second^7,8,9,10,11^; more often, both traits lie downstream of pleiotropic processes, and association data alone will not identify them precisely^9,10^.

Still, pleiotropic processes might be characterized by the SNPs that affect them and by the traits that they affect. In particular, the effect-size covariance matrix for each process specifies the traits that it affects, the directions of those effects, and the SNPs whose association data fit that pattern. If the processes are statistically independent, then the genome-wide genetic covariance matrix is the sum of these component matrices, and pleiotropic decomposition aims to recover them.

Based on the genome-wide covariance alone, many decompositions are equally likely. However, a mechanistically informative decomposition should be *parsimonious*, such that most SNPs belong to zero or one components. Under an unparsimonious model, pleiotropic variants might affect as many processes as they do traits, and little insight is gained. Under a mechanistically informative model, they belong to a single component, and their pleiotropic effects are parsimoniously explained.

Two methods have been proposed that decompose the genetic correlation matrix into components or factors, not exploiting parsimony. Genomic SEM^12^ estimates a factor model matching the observed correlations; this approach can be useful for clustering related traits. Another approach is to use singular value decomposition of the summary statistics matrix, which is equivalent to eigendecomposition of the summary statistics covariance matrix^13,14^. A limitation of both methods is that they operate on the genetic correlation matrix alone, not modeling the effect sizes of individual SNPs, and this approach is unable to exploit parsimony. This limitation is addressed by *mash*^15^, which learns the weights of a gaussian mixture model with components chosen using heuristics. *mash* has been applied to eQTL data across tissues^15^ but not to complex trait association data.

A different approach is to cluster genome-wide significant SNPs. Udler et al.^16^ applied nonnegative matrix factorization to genome-wide significant SNPs for T2D, clustering T2D SNPs by their effects on related phenotypes, and related approaches have also been used elsewhere^17,18^. Julienne et al.^19^ fit a Gaussian mixture model to significant SNPs that passed any of several multi-trait association tests. These methods can provide mechanistic insights into the loci they analyze, but a limitation is that genome-wide significant loci explain a small fraction of heritability for most diseases.

Besides providing mechanistic insights, multi-trait association methods can be used to boost association power. Often, disease association studies are less well powered than those for correlated quantitative traits, motivating cross-trait association methods. MTAG^20^ performs cross-trait meta-analysis with optimal linear weights calculated from the genetic correlation matrix, and a closely related method is HIPO^21^. Other methods include ASSET^22^, which aims to identify association signals present in a subset of traits; PLEIO^18^, which tests the null hypothesis of no association with any trait; CLC^23^, which clusters positively correlated traits and tests the null hypothesis of no association with any cluster; flashfm^24^, which learns a prior for the set of traits that are affected by each SNP; and CTPR^25^, which performs regularized regression with a cross-trait penalty function. Pleiotropic decomposition may improve upon these approaches by modeling a different genetic covariance matrix for SNPs in different components.

## Results

### Overview of Methods

We developed pleiotropic decomposition regression (PDR) to model the distribution of effect sizes across several traits, to estimate shared components of heritability, and to identify the underlying genetic variants. Under the PDR model, the vector of effect sizes for one SNP across several traits, ***β***, is decomposed into a sum of *K* independent components, ***γ*_1_**, …, ***γ_K_***:

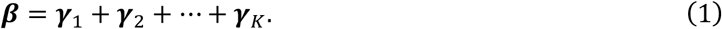

The components might affect different subsets of phenotypes (for example, see Supplementary Figure 1). More specifically, each component ***γ***_*k*_ has a *pattern matrix* Σ_*k*_ parameterizing a scale mixture of multivariate normal distributions:

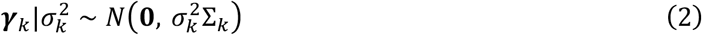

These pattern matrices determine the traits that are affected by each component, as well as the correlation structure across those traits (see Methods for details).

Each SNP has a random vector of *component membership scalars*, 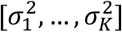, with one entry for each component. The prior distribution over the membership scalars produces parsimony if the smallest scalars have the largest prior probabilities. Conditional on the component membership scalars, the effect-size vector of each SNP follows a multivariate normal distribution, with covariance equal to the component-membership-weighted sum of the pattern matrices.

We are most interested in estimating the genetic covariance explained by each component, which is equal to the pattern matrix of that component times its expected scaling parameter. These are the components of the total genetic correlation matrix:

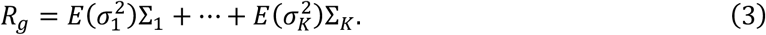

PDR models both trait-specific and pleiotropic components. Trait-specific components have fixed pattern matrices with a single nonzero entry; pleiotropic components have arbitrary pattern matrices that are learned from the data (see Methods).

PDR operates on marginal effect sizes, or associations, as opposed to causal (fine mapped) effect sizes. This choice, common in multi-trait analyses^10,20,6^, is justified because LD alters effect-size magnitudes much more strongly than it does correlations among traits; the genetic correlation matrix is similar for causal and marginal effect sizes (Supplementary Table 1). Moreover, because we model the marginal effect sizes as a sum across components, it is permitted for one associated SNP to tag two causal SNPs belonging to different components. For these reasons, we expect that the proportion of variance explained by a pleiotropic component will be similar to its proportion of heritability explained, and we describe proportions of variance as proportions of heritability below. However, this approach does have important limitations (see Discussion). Effect sizes are measured in per-normalized-genotype units, not in per-allele units.

PDR itself is a method-of-moments estimator in the time domain. We recently developed a time-domain estimator for single-trait effect-size distributions^26^. PDR estimates the empirical characteristic function (ECF) of the association vectors, accounting for GWAS sampling noise and sample overlap, at vector-valued “sampling times”. It estimates the mixing weights by matching the observed ECF with what would be expected under the model using a fast iterative procedure. It updates the pattern matrix of each component using gradient descent. This procedure is repeated with multiple random initializations, the initialization with the best objective function value is selected, and the gradient descent procedure is repeated until convergence. Then, the objective function itself is updated to account for correlations among the sampling times under the estimated model, and the weight estimation and gradient descent procedure is repeated. See Supplementary Figure 2 for a schematic, and see Methods for details on each step.

For each SNP, PDR estimates its component membership scalars, and we can assign SNPs to the component(s) that explain their effects. Integrating over the posterior distribution of these scalars, we compute posterior-mean effect sizes on each trait; these estimates efficiently integrate association data across traits (see Methods).

PDR can be computationally intensive, as it requires a large number of gradient descent steps and random initializations. The parameter space that must be explored grows exponentially with the number of traits and the number of components (Supplementary Figure 3), so it cannot be applied to a large number of traits (here, we analyze 5-6 traits at a time). For a model with six traits, three pleiotropic components, and 10,000 initializations it takes about one hour to run PDR.

Open-source software to run PDR is available (see URLs).

### Simulations

#### Method Comparisons

We compared the performance of PDR against two existing methods, genomic SEM^12^ and singular value decomposition^13^ (called DeGAs in Tanigawa et al., here SVD), which decompose the genetic correlation matrix into factors or components. We simulated data from a model with between one and three “factor-like” components, similar to the factors that these methods estimate (see Methods for a description of the different types of components); this model itself was inferred using PDR from six real metabolic traits (see below; see Supplementary Table 15 for true model parameters). The input to SEM and SVD is a covariance matrix (as SVD applied to the summary statistics matrix is equivalent to eigendecomposition of the sample covariance matrix). Instead of using a noisy estimate based on simulated summary statistics, we used the true genetic correlation matrix as input to the existing methods, illustrating their limiting behavior at large sample size. For PDR, we used the noisy summary statistics as input, with a realistic amount of noise (see Methods and Supplementary Figure 4 for simulation details). We quantified the root sum of squared differences between the simulated and estimated factor weights after reordering and normalization (see Methods). Simulations were performed without LD; the distinction between causal and marginal effect sizes is not central to any of the methods.

Under a model with only one pleiotropic component, the genetic correlation matrix contains most of the information that is needed to infer the factor weights. As a result, all three methods approximately recovered the factor loadings (Figure 1b), with low estimation error (Fig 1a). This simulation corresponds to a situation where all pleiotropic SNPs have a similar pattern of shared effects across traits.

**Figure 1:**
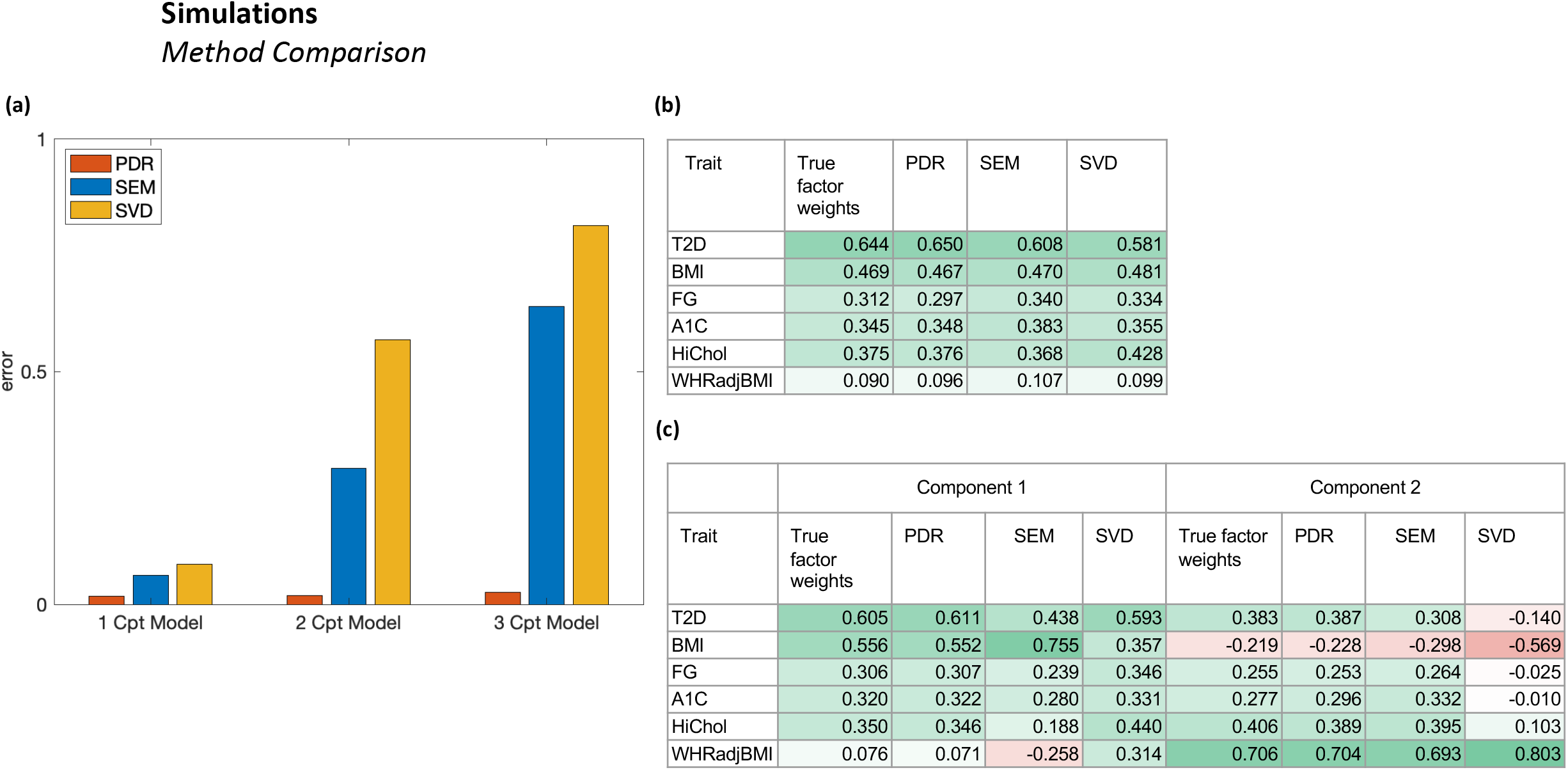
Method comparisons in simulated data. We compared PDR against genomic SEM and SVD in simulations with one, two or three factor-like pleiotropic components. (a) Normalized sum of squared errors between true and estimated factor weights. For PDR, error was averaged across ten replicates. For each method, we re-ordered the factors and re-oriented the positive direction to minimize the error. See Supplementary Table 3a for numerical results. (b) True and estimated factor weights under a model with one pleiotropic component. The cells are color coded from red (−1) to green (1). (c) True and estimated factor weights under a model with two pleiotropic components. In (b-c), the PDR factor weights are from a representative replicate.

We simulated from models with two or three pleiotropic components, where different sets of SNPs have distinct multi-trait association patterns. In these simulations, the genetic correlation matrix does not contain the information that is needed to distinguish the components from each other, so we expected that SEM and SVD would perform less well. Indeed, neither method recovered the components accurately (Figure 1a); SEM partially recovered the two-component model (Figure 1c) but not the three-component model (Supplementary Figure 5), while SVD did not recover the components under either model. PDR produced near-zero error under both models (error=0.02,0.03).

We simulated from models that were designed to favor SEM and SVD respectively (see Supplementary Table 15 for true model parameters). We applied each method to the six real metabolic phenotypes, identified two factors, and simulated from a model utilizing those factors (see Methods). Under the SEM-derived model, SEM and PDR recovered the factors (error=0.03 and 0.05, respectively), but SVD did not (error=0.27). Under the SVD-derived model, SVD and PDR recovered the factors (error=0.10 and 0.02, respectively), and SEM did not (error=0.20) (Supplementary Figure 6). In summary, PDR was the only method that recovered the true pleiotropic components across all simulations.

### Shared heritability components of CAD

We applied PDR to summary statistics for coronary artery disease (CAD) and five related traits: BMI (r_g_=0.28 with CAD), hypertension (r_g_=0.51), high total cholesterol (r_g_=0.62), triglyceride levels (r_g_=0.33), and HDL levels (r_g_=-0.26) (Supplementary Tables 1 and 4). The high total cholesterol phenotype is highly genetically correlated with LDL cholesterol levels (r_g_=0.70), and it has a stronger genetic correlation with CAD (r_g_=0.62 vs r_g_=0.20). We used European-ancestry summary statistics from CARDIoGRAM^27^ for CAD (N=77,210) and from UK Biobank^28,29^ for the other traits (average N=380,413). PDR cannot be applied to summary statistics from different ancestry groups, as it does not model differences in allele frequency or LD across datasets (see Discussion).

We fit models with between one and four pleiotropic components. No restrictions were imposed on their pattern matrices; we also investigated models with factor-like components, which are constrained to have rank one, and determined that they did not fit as well (see Supplementary Table 2 and Methods). A model with two pleiotropic components fit significantly better than a model with one (p=4×10^−50^), and a model with three fit better than a model with two (p=8×10^−5^), but a model with four did not fit better than a model with three (p=1.0) (Supplementary Table 5, Supplementary Figure 7). We subsequently focused on the three-component model. None of its trait-specific components were statistically significant (Supplementary Table 6a), indicating that all SNPs could plausibly be assigned to one of the pleiotropic components.

The three pleiotropic components were closely related to BMI, hypertension, and cholesterol respectively (Figure 2a). The BMI-related component explained most of the heritability of BMI and had strongly correlated effects on all the other traits; it explained about 18% of CAD heritability. (“Heritability” refers to marginal effect size variance; see Overview of Methods.) The hypertension-related component explained about half of the heritability for both hypertension and CAD, but it had little effect on BMI, despite the strong genetic correlation between BMI and hypertension (r_g_=0.38). The cholesterol-related component explained about half of the heritability of high total cholesterol, triglycerides, and HDL, and about 14% of CAD heritability; it had little effect on BMI or hypertension.

**Figure 2:**
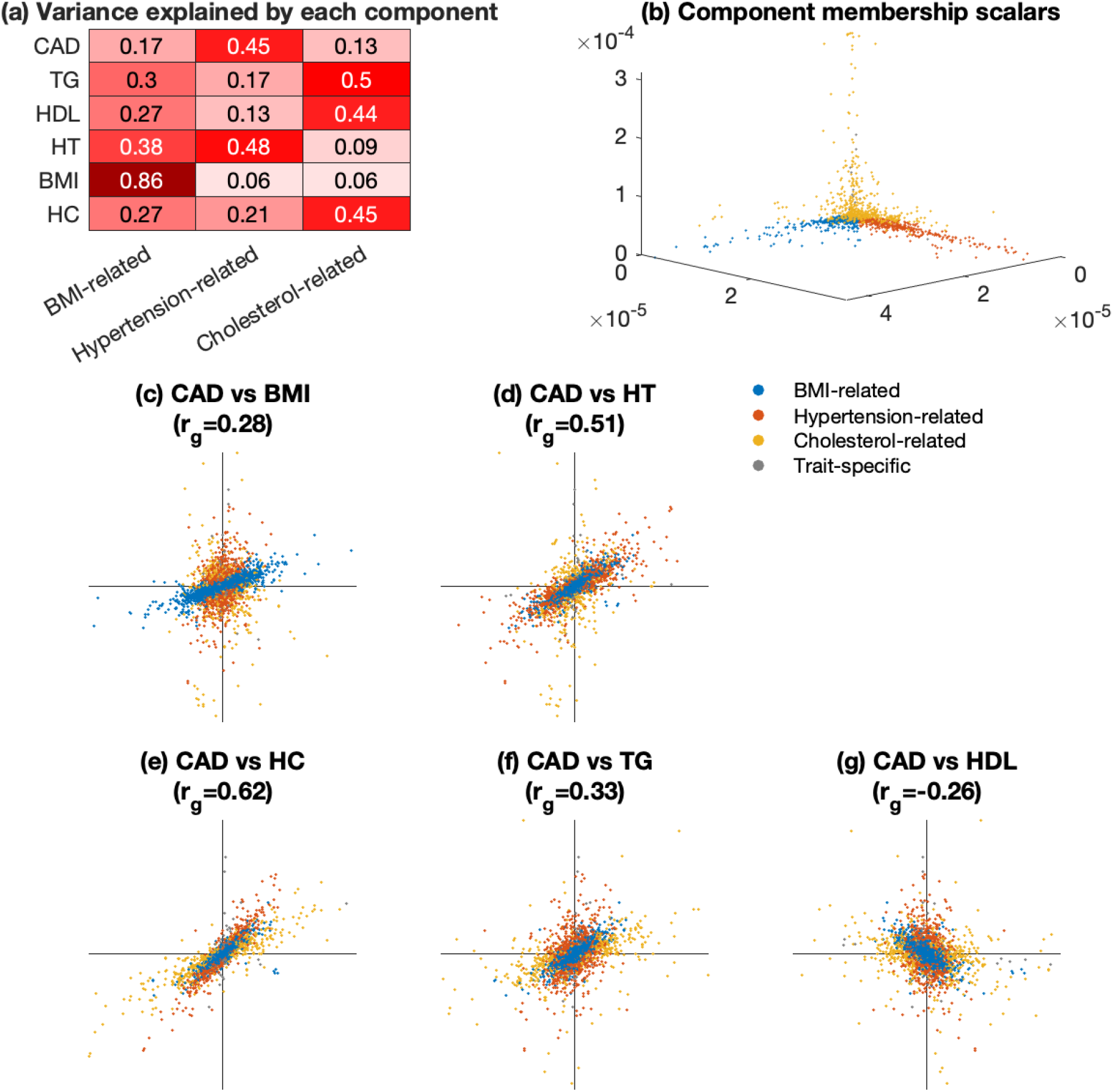
Shared components of heritability for CAD. We applied PDR to coronary artery disease (CAD) and five genetically correlated traits. We performed LD pruning, computed posterior-mean effect sizes of the remaining M=11k SNPs, and assigned each SNP to one component for visualization. (a) Variance explained by the three pleiotropic components for each trait. (b) Posterior-mean component membership scalars for each SNP, colored by component assignment. (c) Posterior-mean effect sizes on CAD (y-axis) and BMI (x-axis); (d) posterior-mean effect sizes on CAD and hypertension; (e) posterior-mean effect sizes on CAD and high total cholesterol; (f) posterior-mean effect sizes on CAD and triglyceride levels; (g) posterior-mean effect sizes on CAD and HDL levels. See Supplementary Tables 10 and 12 for the fitted model parameters and pruned SNP information, respectively.

We computed posterior-mean effect size estimates for each SNP, using the PDR model as a prior (see Methods). These estimates efficiently share information across traits, and we estimate that they explain 40% of the variance in the true CAD effect sizes (r^2^=0.40, s.e.=0.03). In contrast, the original summary statistics are expected to explain a much smaller fraction (r^2^=0.10, s.e.=0.01). We validated our posterior-mean effect size estimates in out-of-sample replication analyses for asthma and T2D (see below).

We used our posterior-mean effect sizes to visualize the estimated effect-size distribution (Figure 2). We performed LD pruning, retaining a set of 11,914 approximately independent SNPs (r^2^<0.01) whose posterior-mean effect sizes were in the top 1% of all SNPs for at least one trait. Most SNPs clustered cleanly into a single component (Figure 2b), and we assigned each SNP to a single component for the purpose of visualization (see Methods).

BMI-related SNPs affect every trait in the cluster, especially hypertension, with highly correlated effect sizes. Despite affecting multiple CAD risk factors, these SNPs mostly have small effects on CAD, and they do not explain a large fraction of its heritability. 80% of SNPs were assigned to this component, reflecting the high polygenicity of BMI. We were concerned that the BMI-related component may be confounded by socioeconomic status, and to assess this, we added educational attainment (EA) (Supplementary Table 4) to the coronary cluster. The estimated components were similar, and there was no change in the interpretation of the BMI component (Supplementary Figure 8).

Whereas BMI-related SNPs also affect hypertension risk, hypertension-related SNPs mostly do not affect BMI. Hypertension-related SNPs have some of the strongest associations with CAD, and they explain a larger fraction of CAD heritability despite the larger number of BMI-related SNPs (Figure 2a). The relative effect size on CAD vs. on hypertension (Figure 2d slope) is similar for hypertension-related and BMI-related SNPs. This observation suggests that the genetic correlation between BMI and CAD might be explained by the effect of BMI-related SNPs on hypertension.

BMI-related and hypertension-related SNPs also affect high total cholesterol, whereas cholesterol-related SNPs mostly do not affect hypertension or BMI (Figure 2c-e). For hypertension-related SNPs, their relative effect on CAD vs. on cholesterol appears to be larger than it is for cholesterol-related SNPs. This difference contrasts with the relationship between BMI and hypertension; it suggests that hypertension-related SNPs, even if they do affect cholesterol levels, have effects on CAD that are not mediated by cholesterol.

All three components affect both triglyceride levels and HDL levels, which are highly negatively correlated (r_g_=-0.58). Triglycerides, but not HDL, are thought to have a causal effect on CAD risk^7,8^. SNPs affecting these traits have heterogenous effects on CAD, and PDR does not strongly distinguish between the genetic architecture of these traits (Figure 2f-g). The effects of cholesterol-related SNPs on CAD are much more strongly correlated with their effects on high total cholesterol (Figure 2e).

We identified the closest gene to each SNP and performed a gene-set enrichment analysis using differentially expressed gene sets that were previously calculated^30^ from GTEx gene expression data^31^ (see Methods and URLs). These enrichment analyses compare genes belonging to one component with genes belonging to others, not with the set of all genes. The BMI-related pleiotropic component was significantly enriched in brain tissues (Supplementary Table 7), consistent with previous findings on BMI^30^. The cholesterol-related component was most strongly enriched in visceral adipose tissue. This component also was nominally enriched in liver tissue, but not after multiple hypothesis correction^32^.

### Shared heritability components of asthma

We applied PDR to UK Biobank summary statistics for asthma and four genetically correlated traits: BMI (r_g_=0.15 with asthma), eczema or allergies (r_g_=0.69), eosinophil count (r_g_=0.45), and FEV1/FVC (r_g_=-0.33) (see Supplementary Table 4 and URLs). The eczema/allergy phenotype does include asthma-like symptoms like wheezing. We selected a model with three pleiotropic components, which fit better than smaller models (Supplementary Table 5, Supplementary Figure 9). Collectively the pleiotropic components explained ~90% of heritability; two trait-specific components, for eczema and eosinophil count, were statistically significant (Supplementary Table 6).

We identified components related to BMI, inflammation, and pulmonary function (Figure 3a), and these components were mostly disjoint (Figure 3b). BMI-related SNPs explained the majority of BMI variance and had correlated effects on asthma, while SNPs belonging to the other components had little or no effect on BMI (Figure 3a,c,d). The pulmonary component associated most strongly with FEV1/FVC, and it had strongly correlated effects on asthma and on eczema, but not on eosinophil count or BMI. The inflammatory component associated strongly with eosinophil count, eczema and asthma, but not with BMI or FEV1/FVC.

**Figure 3:**
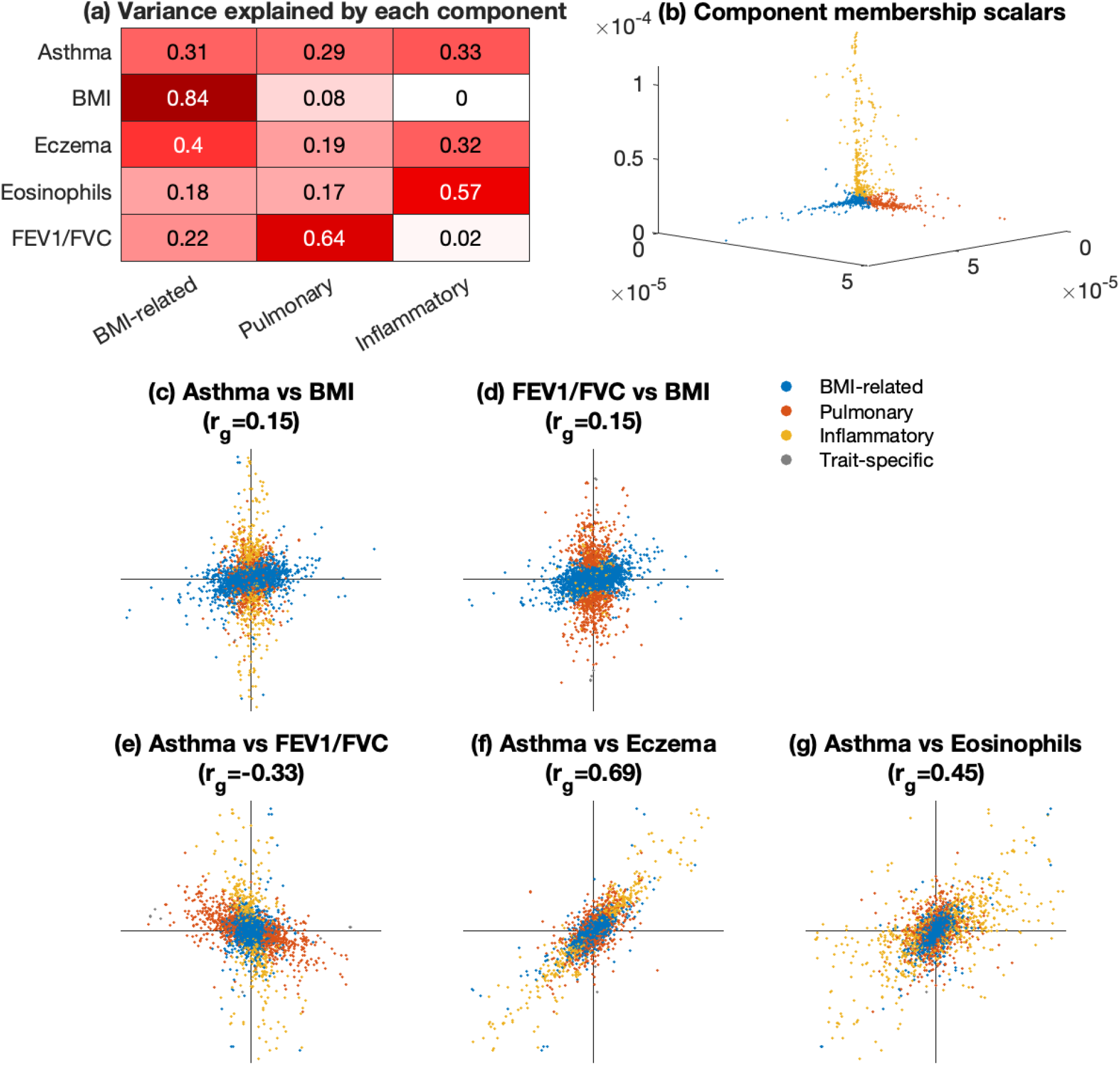
Shared components of heritability for asthma. We applied PDR to asthma and four genetically correlated traits, calculated poster-mean effect sizes for LD pruned SNPs, and assigned SNPs to components. (a) Variance explained by the three pleiotropic components for each trait. (b) Posterior-mean component membership scalars for each SNP, colored by component assignment. (c) Posterior-mean effect sizes on asthma (y-axis) and BMI (x-axis); (d) posterior-mean effect sizes on FEV1/FVC and BMI; (e) posterior-mean effect sizes on asthma and FEV1/FVC; (f) posterior-mean effect sizes on asthma and eczema/allergies; (g) posterior-mean effect sizes on asthma and eosinophil count. See Supplementary Tables 10 and 12 for the fitted model parameters and pruned SNP information, respectively.

These findings align with the hypothesis that asthma has distinct obesity-related and allergy-related subtypes^33^, loosely corresponding to adult-onset and childhood-onset asthma respectively^34^. We additionally identify a third component related to pulmonary function; most SNPs that increase FEV1/FVC are associated with decreased risk of asthma (Figure 3e). This consistent pattern suggests that high baseline pulmonary function protects against asthma or reduces the probability of diagnosis. In contrast, inflammatory SNPs had a strong effect on asthma but not on FEV1/FVC (Figure 3e), which was measured at baseline (not after exercise or exposure to allergens).

BMI, asthma, and FEV1/FVC have seemingly discordant pairwise correlations: BMI is slightly positively correlated with both asthma and FEV1/FVC (rg = 0.15 and 0.15), but FEV1/FVC and asthma are negatively correlated (rg = −0.33). PDR was able to explain these nontransitive correlations by uncovering distinct underlying sets of SNPs. BMI-related SNPs drive the positive correlations with BMI; pulmonary SNPs, which do not affect BMI, drive the negative correlation between FEV1/FVC and asthma (Figure 3c-e).

We performed gene-set enrichment analyses and identified 7 significant component-tissue pairs (Supplementary Table 8). Like the BMI-related component of the CAD cluster, the asthma BMI-related component was significantly enriched in brain tissues. The inflammatory component was enriched in the spleen and small intestine, and the pulmonary component was enriched in the esophagus and the tibial artery.

We also applied PDR to a cluster of metabolic traits consisting of type II diabetes (T2D) and five genetically correlated traits: BMI (r_g_=0.56 with T2D), fasting glucose (r_g_=0.63), HbA1c (r_g_=0.67), high total cholesterol (r_g_=0.46), and waist-hip ratio adjusted for BMI (WHRadj) (r_g_=0.27) (Supplementary Figures 10-11). We identified components related to BMI, waist-hip ratio, and glucose/A1c. We investigated possible collider bias between BMI and WHRadj^35^ and determined that confounding was unlikely (Supplementary Figure 12).

### Out-of-sample validation

We obtained non-UK Biobank summary statistics for T2D^36^ and asthma^37^ in order to validate our posterior-mean effect size estimates (Supplementary Table 4). We defined the *replication r^2^* as the squared correlation between true and estimated effect sizes; it quantifies the fraction of variance in the true effect sizes explained by the estimated effect sizes. For subsets of SNPs, we estimated the *per-SNP variance explained*, which is the replication r^2^ for those SNPs times their variance enrichment (see Methods). Both quantities can be predicted without the use of replication data by taking the sample variance of the posterior-mean effect sizes. These predictions are only accurate if the posterior-mean effect sizes are unbiased, and we validate these predictions in the non-UK Biobank replication data. We also calculated the replication r^2^ of the original summary statistics, and we normalized the validation data such that the predicted and observed replication r^2^ for the original summary statistics were equal (see Methods).

For asthma, PDR posterior-mean effect sizes had a predicted replication r^2^ of 0.59 (s.e.=0.06) and an observed replication r^2^ of 0.66 (0.11) (Figure 4a). The latter represents a 94% improvement over the original summary statistics (r^2^=0.34, s.e=0.005). For T2D, PDR had a predicted and observed replication r^2^ of 0.47 (s.e.=0.02) and 0.46 (0.04) respectively (Figure 4b). Again, these numbers represented a large improvement, of 70%, over the original summary statistics (r^2^=0.27, s.e.=0.003).

**Figure 4:**
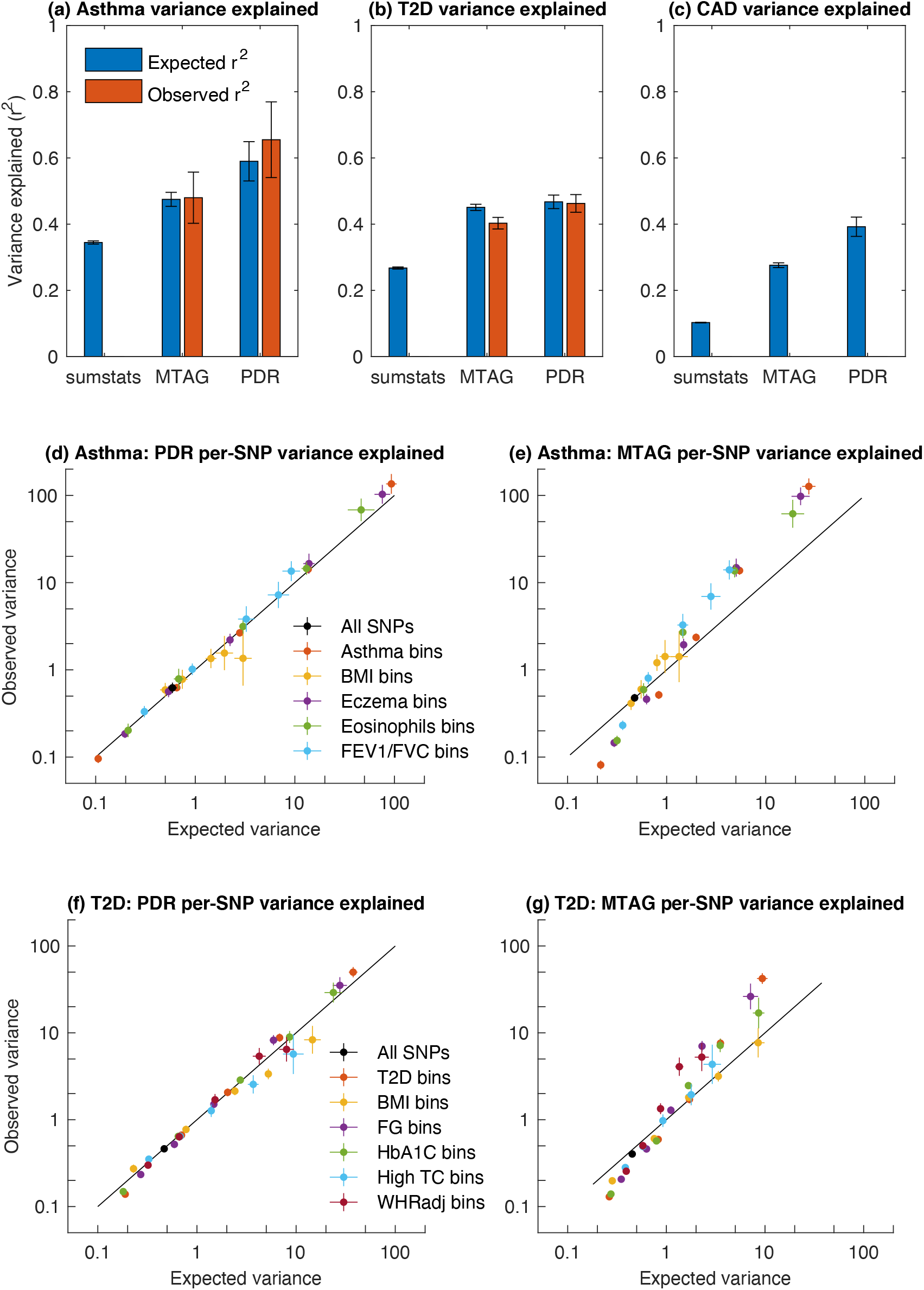
Out-of-sample replication of posterior-mean effect sizes. (a-c) Replication r^2^ for the original summary statistics, MTAG/BLUP, and PDR. Expected r^2^ is calculated within the training sample, and observed r^2^ is calculated out of sample (for the original summary statistics, expected and observed r^2^ are equal due to normalization). For CAD, replication data were unavailable, so only the expected r^2^ is reported. Error bars indicate one standard error (calculated via block jackknife). (d-e) We binned SNPs by their estimated effect size (*χ*^2^ statistic) on each trait in the asthma cluster and computed expected and observed variance explained in the effect size of these SNPs on asthma, using either PDR (d) or BLUP (e). When expected variance is greater than observed, it indicates that effect sizes are overestimated, and vice versa. (f-g) Expected and observed variance explained for T2D. See Supplementary Table 13 for numerical results.

For CAD, replication data were unavailable, but the expected r^2^ was much larger for PDR (0.39, s.e.=0.03) than for the original summary statistics (0.10, s.e.=0.001). This ~300% improvement, although not confirmed with replication data, was much larger than what we observed for T2D and asthma. The CARDIoGRAM association data started at a lower baseline (r^2^ = 0.10 vs 0.27 and 0.34), and non-UK Biobank summary statistics might derive greater benefit from being pooled with UK Biobank data.

We compared PDR posterior-mean effect sizes with estimates obtained using MTAG, which performs linear meta-analysis using optimal weights^20^. MTAG estimates are proportional to the posterior-mean effect sizes that would be obtained with a multivariate normal prior; applied to phenotype prediction, this approach is commonly called the best linear unbiased predictor, or BLUP^38,39^. Because they are proportional, MTAG and BLUP have identical replication r^2^. BLUP/MTAG had replication r^2^ about midway between PDR and the original summary statistics (Figure 4a-c).

For both PDR and BLUP, predicted and observed replication r^2^ were approximately concordant, indicating that their posterior-mean estimates were unbiased on average:

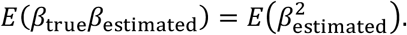

This type of unbiasedness is related to the Bayesian unbiasedness of Goddard et al.^39^, that

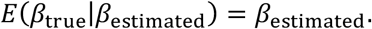

However, the latter definition of unbiasedness is stronger, as posterior-mean effect sizes can be biased for SNPs with large or small effect sizes in such a way that the biases cancel out on average. Well-calibrated posterior-mean effect sizes should be unbiased on average not only for the set of all SNPs, but for any subset of SNPs binned by their estimated effect sizes on each trait (because binning is like conditioning). Any model misspecification is expected to produce bias. For example, if PDR overestimated the fraction of BMI-related SNPs that affected asthma, then its posterior-mean effect sizes on asthma would be upwardly biased for BMI SNPs, and the predicted variance explained would be greater than observed.

We compared the expected and observed variance explained for subsets of SNPs binned by their *χ*^2^ statistics for each trait (Figure 4d-g). PDR predicted and observed variance explained were approximately concordant for every subset of SNPs, both for asthma (Figure 4d) and for T2D (Figure 4f). Across bins, both the expected and observed values spanned more than two orders of magnitude, which is roughly the range of effect sizes that is required to explain most complex-trait heritability^26^. This concordance indicates that PDR produces approximately unbiased posterior-mean effect sizes for SNPs across the entire range of the multi-trait effect size distribution.

In contrast, MTAG/BLUP posterior-mean effect sizes exhibited bias (Figure 4e,g). For small-effect SNPs, their effect sizes were overestimated; most of these SNPs are probably null. For large-effect SNPs, their effects were underestimated, as a multivariate normal (infinitesimal) prior does not allow for a small percentage of SNPs with very large effect sizes. We note that the MTAG estimator is not described by Turley et al.^20^ as a posterior-mean estimator, even though it is closely related to the BLUP posterior-mean estimator. An advantage of its linear meta-analysis approach is that the resulting effect-size estimates have similar sampling properties as the original summary statistics, whereas the sampling distribution of our posterior-mean estimates is more difficult to characterize.

## Discussion

We developed pleiotropic decomposition regression (PDR) to infer shared components of heritability across traits, to characterize their pleiotropic mechanisms, and to identify their underlying genetic variants. We applied PDR to genetically correlated trait clusters associated with CAD, asthma, and T2D. The components that were identified corresponded to established risk factors for these diseases, but unlike the risk factors themselves, the components had mostly distinct underlying variants. In out-of-sample validation analyses, we found that PDR posterior-mean effect size estimates were substantially more accurate than the original GWAS estimates, and that they were unbiased, indicating that the PDR model does fit the data.

The components identified by PDR most likely represent heterogenous clusters of mechanisms, not specific pathways or intermediate phenotypes. We did not observe that specific pathways, which would presumably correspond to small sets of loci with homogenous patterns of association across traits, could be identified by PDR. A measure of homogeneity within a component is the strength of its correlations; strong correlations indicate that most SNPs within the component have similar relative effect sizes on each trait. Most of our inferred components had relatively weak correlations. These weaker correlations could reflect limitations of the PDR method or limited power in the GWAS data; alternatively, they might reflect the limited mechanistic resolution of common-variant associations.

Our posterior-mean effect sizes have greatly increased replication r^2^ compared with the original summary statistics, and this increase in estimation accuracy suggests that PDR might be used to discover new loci. However, PDR does not produce p-values for individual SNPs, and it is not clear what metric should be used, or what threshold, to determine significance. One approach would be to use the not-by-chance true positive rate (NTPR^26^), which can be calculated using PDR, but a critical limitation is that PDR does not model LD among nearby SNPs. The NTPR of lead SNPs, not in LD with any stronger association, is expected to be higher than that of tag SNPs with similar *χ*^2^ statistics, as the true-positive rate of tag SNPs is bolstered by their more strongly associated LD partners.

Increased replication r^2^ could also be useful for polygenic risk prediction, and several existing PRS methods do incorporate multi-trait data^25,40,41,42^. Explicitly modeling multi-trait effect size distributions could be a promising approach, and our results highlight CAD in particular as a trait that would benefit (Figure 4c), but again a limitation of using PDR itself is that it does not model LD. Methods that explicitly model LD patterns generally outperform models that use simple LD pruning, although not by large margins^43,44^. PDR posterior-mean estimates could easily be paired with a simple LD pruning heuristic, and we do not know whether the advantages of explicitly modeling multi-trait effect size distributions would outweigh the limitations of LD pruning.

PDR is not applicable to datasets from different ancestries, again because it does not currently model LD. Cross-population genetic correlations can be estimated using methods that explicitly model differences in LD patterns and allele frequencies among different studies^45,46^. It is essential to account for these differences, because otherwise the observed genetic architecture will seem to be much more divergent across populations than it actually is^47^.

Besides the fact that it does not model LD, PDR has other important limitations. First, it cannot be applied to more than around six traits due to its computational cost. Because of this limitation, users must decide before applying PDR which traits are most interesting to analyze, and we recommend choosing them based on their genetic correlations. Second, PDR does not currently produce standard errors, and the typical approach of using block-jackknife standard errors is computationally intensive. It may be possible to estimate standard errors using second derivatives of the objective function instead, but we have not validated this approach. PDR does produce p-values for its components, so users can be confident that the estimated components are not spurious. Despite these limitations, PDR is a sophisticated estimator of the multi-trait effect size distribution that can be used to produce biological insights and to improve association accuracy.

## Acknowledgements

This research was supported by a grant from the Simons Foundation (704413). We are grateful to Alex Bloemendal, Eric S. Lander, Ajay Nadig, Ben Neale, Alkes L. Price, Dan Weiner, and Kenneth Westerman for helpful discussions, comments, and suggestions.

## URLs

UK Biobank summary statistics: https://alkesgroup.broadinstitute.org/sumstats_formatted/;

1000 Genomes LD scores: https://alkesgroup.broadinstitute.org/LDSCORE/;

Trans-National Asthma Genetic Consortium (TAGC) asthma summary statistics: http://ftp.ebi.ac.uk/pub/databases/gwas/summary_statistics/GCST006001-GCST007000/GCST006862/

CARDIoGRAM coronary artery disease summary statistics: http://www.cardiogramplusc4d.org/data-downloads/

GIANT summary statistics: https://portals.broadinstitute.org/collaboration/giant/index.php/GIANT_consortium_data_files

Open-source software for PDR: https://github.com/jballard28/PDR;

MTAG: https://github.com/JonJala/mtag

Genomic SEM: https://github.com/GenomicSEM/GenomicSEM

PLINK 2.0: www.cog-genomics.org/plink/2.0/

acmeR: https://cran.r-project.org/package=acmeR

## Methods

### PDR Model

PDR fits a model for the distribution of marginal effect sizes (associations, inclusive of LD) across several traits. It models this vector, ***β***, as a sum of *K* independent random vectors, ***γ***_1_, …, ***γ***_*K*_, corresponding to the *K* components. If there are three traits:

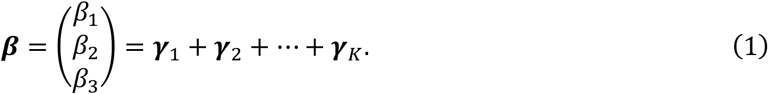

For each component, its random effect-size vector follows a scale mixture of normal distributions. This mixture is parameterized by a *pattern matrix* Σ_*k*_, which determines which traits are affected as well as the pattern of correlations across traits:

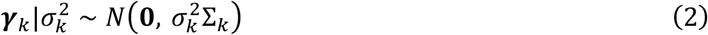

The random variable 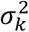, which differs across SNPs (unlike Σ_*k*_), describes component membership, and we call it the component membership scalar. It follows a categorical distribution, leading to a discrete mixture distribution for ***γ***_*k*_:

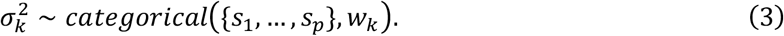

*s*_1_, …, *s*_*p*_ are the possible values of *σ*^2^, and they are fixed, rather than being learned from the data; zero is one of these values, which allows for SNPs that have no effect on the component. *w*_*k*_ is the vector of mixing weights, and it is learned from the data; it describes what fraction of SNPs have a large effect, a small effect, and no effect on the component.

The pleiotropic pattern matrix is an interpretable parameter. For example, the matrix

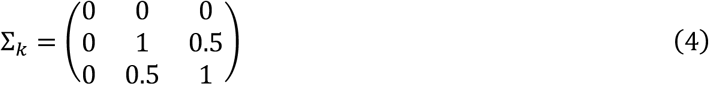

represents a component that affects traits two and three, with positively correlated effect sizes, but not trait one. We report estimates of the variance explained by each component, which is proportional to Σ_*k*_; in particular, the variance explained is:

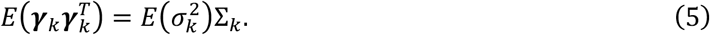

Because the components are independent, the total genetic covariance is

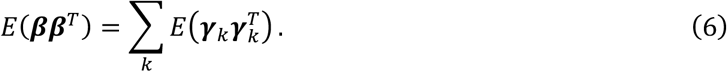

A scale mixture of multivariate normal distributions was previously used by Urbut et al.^1^ in their method *mash*, which they applied to cross-tissue eQTL data. For *mash*, the primary estimation task is to estimate the mixture weights, whereas for PDR, the primary task is to estimate the pattern matrices. *mash* does include “model-driven” covariance matrices, but these are constructed using a heuristic pre-processing step. For applications to complex traits, it is essential to learn the pattern matrices directly from the data because these matrices, not the mixing weights, are the biologically interpretable model output. A second difference is that PDR models a convolution (sum) of components, rather than a mixture; this choice allows a single SNP to affect multiple components, which is desirable especially because we are modeling the distribution of marginal effect sizes. It is possible to use PDR with a mixture-of-components model if desired.

In practice, we are often investigating the genetic basis of a focal trait, like coronary artery disease (CAD), by means of its risk factors. PDR does not distinguish between a focal trait and its risk factors or supporting traits; in particular, there is no guarantee that all of the components it identifies will actually affect the trait of interest. However, traits can be selected that are likely to be informative for a focal trait, and results can be interpreted with a focal trait in mind.

### Types of components

The components ***γ***_*k*_ can be either trait specific or pleiotropic. If component one is trait one specific, its pattern matrix has only one nonzero entry:

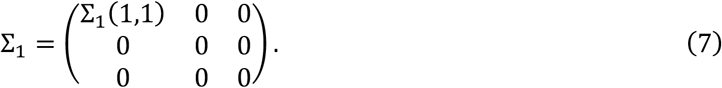

It has one free parameter, corresponding to this nonzero entry. We always include trait-specific components, so that we do not spuriously assign too much heritability to a pleiotropic component. In practice, however, these often do not improve the model fit; we report p-values for whether the model including these components fits better than one without them (see below).

For a pleiotropic component, its covariance matrix can be any positive semidefinite matrix, and it is parameterized as:

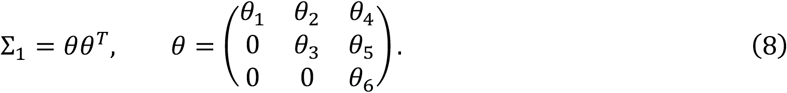

That is, it is parameterized by its Cholesky decomposition, and we learn the Cholesky factors. This choice ensures that Σ is positive semidefinite without having to explicitly constrain it.

PDR can also learn “factor-like” pleiotropic components, which are less general than the generic (“full-rank”) pleiotropic components; we utilize these components for the purpose of method comparisons (see below). Factor-like components are parameterized as:

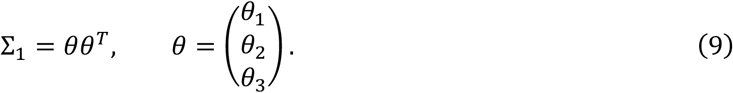

If the components were allowed to be statistically dependent, then it would be possible to model ***β*** using *only* trait-specific components, and they would inherit the same dependency structure as the entries of ***β***. Therefore, we require that the components are statistically independent, forcing the statistical dependencies among traits to be captured by pleiotropic components.

We briefly note some common concerns that are not relevant to PDR or the PDR model. First, we ignore LD, only modeling the distribution of *marginal* effect sizes (inclusive of LD). This choice is justified because LD is expected to affect the magnitude of effect-size vectors, rather than their angles (see Supplementary Table 1). Second, PDR is equally applicable to continuous and dichotomous traits, as the specific units of the effect sizes (e.g. centimeters or log-odds) is unimportant. Third, pleiotropy is sometimes confused with epistasis, a completely different multivariate phenomenon; we model only additive effects.

### Time-domain moment equations

PDR optimizes a feasible generalized least squares (FGLS) objective function with time-domain moment equations. The objective function is:

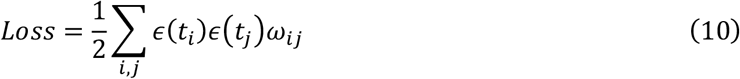

where *t*_*k*_ is the *k*th vector-valued “sampling time”, and *ε*(*t*) is the difference between the expected and empirical characteristic functions evaluated at *t*. *w*_*ij*_ is element (*i*, *j*) of a moment equation precision matrix (inverse-covariance matrix) which is itself estimated from the data (see below).

The characteristic function (CF) of the model is a function of the sampling time *t*:

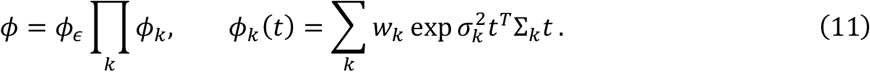

*ϕ*_ε_ is the CF of the noise term. We assume that the sampling errors follow a multivariate normal distribution with covariance equal to the cross-trait LD score intercept matrix, as justified elsewhere^2,3^. With cross-trait LD score intercept covariance matrix Σ_ε_, the noise CF is:

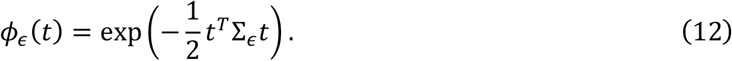

PDR matches the model CF with the empirical characteristic function (ECF), which is a function of the data:

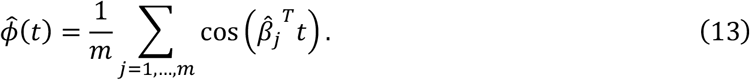

The cosine is the real part of the ECF; the imaginary part is zero, because the distribution of 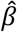 is even 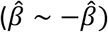. *m* is the number of SNPs. The ECF is only computed once, and subsequent optimizations do not need to iterate over SNPs.

### Sampling times

PDR has one moment equation for each of several “sampling times.” These are vectors of length equal to the number of traits, and they are chosen using a heuristic. The characteristic function (eq. 13) involves the dot product of the sampling times and the effect-size vectors; intuitively, the sampling time *t* provides information about the number of SNPs *j* with different values of 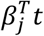. Similar sampling times will provide redundant information.

We sample the sampling times at random on a grid. The grid values are determined by the component scalars: if the possible scalars are {0, *s*_2_, …, *s*_*k*_}, then the grid values are 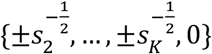. We sample 1000 sampling times at random from the number-of-traits dimensional grid with these sampling times.

### Iterative mixing weight estimation

To estimate the mixing weights, we use an iterative approach that involves performing linear regression at each step. There is a weights vector, *w_k_*, for each component *k*; its length is the number of scalars for that component, where the scalars are *s*_1_, …, *s_p_*. The optimization has the form:

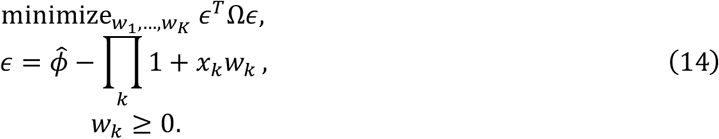

The vector *x*_*k*_, as well as the scalars 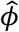 and *ε*, is implicitly a function of time, and it is evaluated at each of the sampling times:

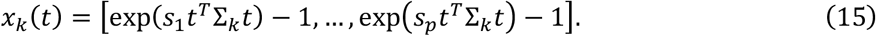

Ω is a regression weights matrix with an entry for each pair of sampling times (see below).

This closely resembles a linear regression problem of 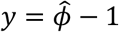 on *X* = [*x*_1_, …, *x*_*K*_]. It can be solved by iterative application of weighted linear regression. We initialize *w*, and then for each component *k*, we regress

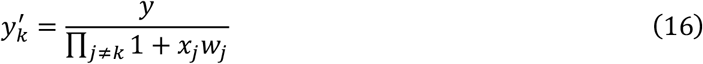

on *x*_*k*_, updating *w*_*k*_ appropriately. The regression is constrained to produce nonnegative coefficients, which has almost no additional computational cost, as previously described^3^. The optimization is convex, and it converges quickly.

### Gradient descent

The pattern matrices themselves are estimated via gradient descent. At each step, we compute the residuals 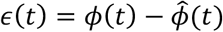, and we compute the gradient of *ϕ*_*k*_(*t*) with respect to the parameters *θ*_*k*_. We choose a step size, take a step along the gradient, re-compute the mixing weights, and repeat until convergence.

The gradient is:

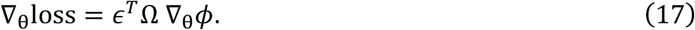

The gradient descent step size is computed via backtracking line search^4^. The search is initialized at twice the previous step size; at each iteration of the search, the step is accepted if

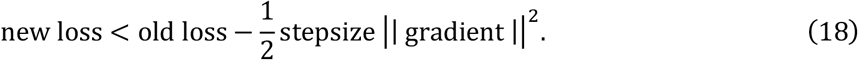

If the condition fails, we reduce the step size by half and try again.

In practice, it can take thousands of steps for the gradient decent procedure to converge. By default, PDR stops after twice failing to improve the loss by a fraction of more than 10^−6^.

### Initialization for gradient descent

The PDR objective function is nonconvex, and multiple random initializations are required in order for it to find the global optimum. We perform a large number of initializations with only 3 gradient descent steps at each initialization, choose the best initialization out of those based on the objective function value, and proceed with further gradient descent steps. PDR has several options for how to choose the initial parameters:

- By default, PDR initializes using the ‘covmat’ method, which initializes the pattern matrix to a random value chosen to be similar to the total genetic covariance matrix (or any other specified covariance matrix). It draws *n* samples from a multivariate normal distribution with the specified covariance (and zero mean), and the sample covariance of these samples is chosen as the initial pattern matrix. The default value of *n* is the number of traits.
- The ‘random’ method initializes the component parameters to random values drawn from the standard multivariate normal distribution.
- The ‘orig’ method initializes the model using the parameter values currently stored in the model. If the parameters of the true model are known, one could initialize to the truth by copying the true model’s components into the model that is being fit and using this initialization method. Since there is no randomization involved in this method, it performs only a single initialization.
- The ‘orig_rand’ method does the first initialization at the parameter values currently set in the model (same as ‘orig’), and then performs the rest of the initializations by drawing random values from a normal distribution based on the data covariance matrix (same as ‘covmat’).
- The ‘near_orig’ method randomly initializes the parameters to values that are within some user-specified distance from the values originally stored in the model. This could be used to explore the parameter space near a specific set of values. For example, it might be of interest to see in simulations if a model initialized near the truth always converges to the true parameters.
- The ‘orig_subset’ method initializes a user-specified subset of components to the initial parameter values (similar to ‘orig’) while randomizing the other components (similar to ‘random’).

PDR can be computationally intensive. Although each gradient descent step is fast, a large number of steps (100-10,000) are usually required for convergence. Moreover, the optimization is nonconvex, and many random initializations (~1,000, with only a few steps per initialization) are required in order to be confident that the global optimum is found. We evaluated in simulations the probability of identifying the global optimum as a function of the number of initializations (Supplementary Figure 13). The cost scales with the number of traits and the number of components, but not with the number of SNPs or the number of individuals in the GWAS. A straightforward maximum-likelihood or pseudolikelihood estimator, evaluating the likelihood at each SNP, would be infeasible. To some extent, the cost is unavoidable, because the space of models that must be explored grows exponentially with the number of traits. Nonetheless, it is a limitation of PDR that it cannot feasibly be applied to more than about ten traits.

### Regression weights

PDR is a feasible generalized least squares estimator (FGLS), which means that it determines its own regression weights based on the data. The motivation for this approach is that if the true effect-size distribution is known, then it determines the sampling variance of the residuals; in particular, if there is no LD, then

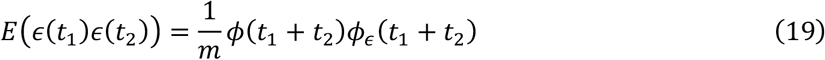

where *ϕ* is the true CF. The optimal regression weights matrix is the precision matrix, Ω, or the inverse of the covariance matrix. We iteratively update this matrix during the model fitting procedure, adapting the covariance to match the current estimate of the model parameters; by default, PDR performs twenty updates to this matrix, which is sufficient for convergence. We handle LD by using a block-jackknife with *m* = 100 blocks, assuming independent jackknife blocks instead of assuming independent SNPs; this approach is commonly used to compute standard errors^5^.

The regression weights we calculate can potentially lead to instability when a collection of sampling times have highly correlated ECF values (which occurs when the sampling times themselves are highly similar). Instead of attempting to remove these correlations, it can be better to simply prune them, resulting in a smaller set of linear combinations of sampling times. This is done using a tolerance parameter; we project the ECF onto the lower-dimensional subspace spanned by the singular vectors whose singular values are greater than this tolerance. In practice we found that this is important to obtain well-calibrated p-values, but it has little effect on the point estimates. We calculate p-values for model comparisons using a tolerance parameter of 0.005; point estimates are computed with a tolerance of zero.

Another potential issue is that occasionally, the model will oscillate between different weights matrices and optima, leading to slow convergence. This issue is addressed with a dampening factor *d* ∈ [0,1): when updating the weights matrix, we update it to Ω_1_ = *d*Ω_0_ + (1 − *d*)Ω, where Ω_0_ is the previous weights matrix and Ω is the new weights matrix. We use *d* = 0.5 by default. This parameter affects the rate of convergence, but not the eventual solution.

### Hypothesis testing

We perform hypothesis testing to compare nested models with different numbers of components. First, we fit the null (smaller) model, and we compute the covariance of the residuals that would be expected if that model were true. Then, we fit the alternative model using that same covariance matrix, we compute the loss 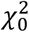 and 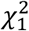 of the null and alternative respectively, and we compute the F-statistic:

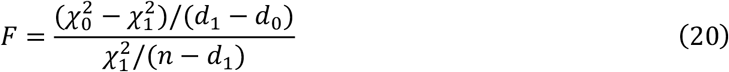

where *n* is the rank of the weights matrix Ω, and *d*_0_ and *d*_1_ are the number of parameters in the null and alternative models respectively. Under the null, this statistic follows an F test with degrees of freedom equal to *d*_1_ − *d*_0_ and *n* − *d*_1_.

### Posterior-mean effect sizes

Using the distribution of effect sizes estimated by PDR, we estimate the posterior-mean effect sizes by calculating 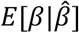, where *β* is the true effect size and 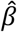 is the effect size from the original summary statistics. Taking advantage of the fact that the Gaussian distribution is conjugate to itself, the mean of the posterior distribution has a closed-form solution; however, it does require us to sum over the posterior distribution of the scaling parameters. In principle, this might be computationally demanding, because there are exponentially many possible scaling parameter assignments for each SNP. In practice, we set a threshold and only examine combinations of scaling parameters whose prior probability is greater than that threshold; this approach is expected to be successful, because most combinations of scaling parameters are extremely unlikely. This threshold is set at 10^−6^.

We integrate over sufficiently likely values:

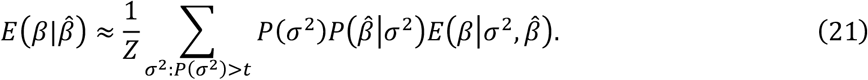

In this expression, *P*(*σ*^2^) is the product of the appropriate mixture weights, 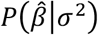 is the Gaussian likelihood of 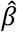 with covariance matrix Σ_*ε*_ + Σ_²_ where 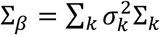, and 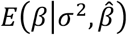 is the Gaussian posterior mean:

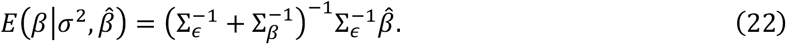

The normalizing constant *Z* is

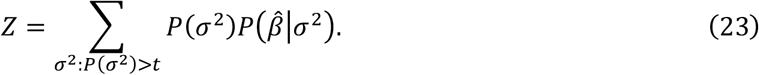

We also computed posterior-mean scaling parameters:

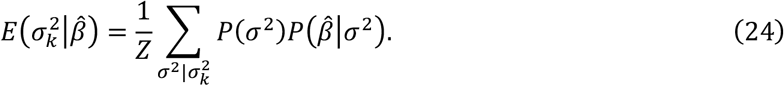

### Method Comparisons

In order to compare the performance of PDR to that of genomic SEM and singular value decomposition (SVD), we simulated data from a model that was itself inferred by applying PDR to real traits. We applied PDR to the metabolic trait cluster (Supplementary Table 4), and we fit models with one, two, and three “factor-like” pleiotropic components (Supplementary Table 15). These factor-like components were constrained to have rank-one covariance matrices; they were designed to match the type of structure that can be inferred by genomic SEM and SVD. Genomic SEM and SVD use a covariance matrix as their input, and we gave them the true covariance matrices which were used to simulate the data.

We ran genomic SEM using the GenomicSEM package developed by Grotzinger et al. (see URLs). We produced an LDSC output object whose covariance matrix was equal to the true covariance matrix, whose sampling covariance matrix was a diagonal matrix with diagonals equal to 0.001, and whose matrix of LDSC intercepts was a zero matrix. The number of SNPs was equal to 10^5^, the number of SNPs in the simulated data. We then ran exploratory factor analysis on this object, setting the number of factors equal to the number of pleiotropic components in the corresponding true model. The factor loadings were taken as the estimated factor weights.

To get the SVD estimates, we performed eigendecomposition on the true genetic correlation matrix. The estimated factor weights were taken to be the top k principal components corresponding to the largest k singular values, where k was the number of pleiotropic components in the true model.

For PDR, we simulated data from each of the true models ten times and ran PDR on the simulated data using the same number of pleiotropic components as the true models. These models were initialized first at the truth, followed by randomly sampling several times from a distribution based on the data covariance matrix (using the ‘orig_rand’ initialization function). This was equivalent to approaching the limit of infinite initializations, which would include the true model parameters; with enough initializations, PDR consistently finds a solution that is close to the truth (Supplementary Figure 13). The factor weights were taken to be the parameter, ***θ*_*k*_**, belonging to each component; the component pattern matrices were 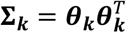.

### Factor Weight Estimation Error

For all three methods, we normalized the factor weight vectors to have unit length to facilitate comparison. The previously outlined procedure produced the estimated factor weights, denoted here as *θ*, but their ordering with respect to the true model was arbitrary. To optimally match the estimated factor weights to the true factor weights, we iterated through each possible way of pairing components between the true and estimated models, while also negating one of the factor weight vectors since the signs are arbitrary in determining the component covariance matrix (***θθ*^*T*^ = −*θ*(−*θ*)^*T*^**), and we chose the factor weight assignments and signs that minimized the total root sum-of-squared differences across all components. Estimation error was quantified as:

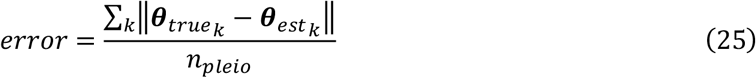

where *n*_*pleio*_ was the number of pleiotropic components.

### Deriving models based on genomic SEM and SVD

We performed method comparisons using models with two pleiotropic components based on factor weights estimated using genomic SEM and SVD (Supplementary Table 15). We first generated genomic SEM and SVD factor weights by applying each method to real data from the metabolic cluster. Then, we modified the two-component PDR model (with factor-like components, estimated as described above), setting the *θ* parameters for each component to the genomic SEM or SVD-derived factor weights. We then modified the weights of the components so that the sum of squared thetas remained unchanged from the original PDR model, thus keeping the total pleiotropic heritability the same within each component. For one of the components in the genomic SEM-derived model, we divided the weights by two so the total heritability did not exceed 1. We then calculated the trait-specific heritability by subtracting the total heritability explained by the pleiotropic components from 1, and then scaled the weights of each trait-specific component so the trait-specific heritability of that component matched this heritability. Using these genomic SEM and SVD-derived models as the truth, we used the same procedure outlined above to estimate the thetas using each of the three methods.

### LD pruning and component assignment

For the purpose of visualization, we performed LD pruning followed by component membership assignment. We estimated posterior-mean effect sizes for each SNP, ranked the top 1% of SNPs for each trait, and sorted SNPs by their topmost rank across traits (for example, a SNP that was ranked first for trait one and third for trait two was retained over a SNP that was ranked second for both traits). We LD pruned this list in a greedy manner, retaining top-ranking SNPs (with larger effect sizes) and discarding SNPs that were in LD with them (r^2^>0.01). We used LD matrices calculated from European samples in the 1000 Genomes Project^6^ using PLINK2^7^ (see URLs).

We assigned the remaining 11,914 SNPs to components using their posterior-mean component membership scalars (see above). For each SNP, and for each component *k*, the expected magnitude of ***γ***_*k*_ is:

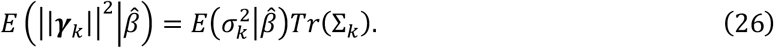

SNPs were assigned to the component *k* with the largest value of 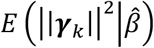.

### Gene-set enrichment analysis for tissue-specific enrichments

We performed a tissue-specific enrichment analysis for each component. The tissue-specific gene sets were previously derived from the GTEx gene expression data^8^ by Finucane et al.^9^ (see URLs). Using the LD pruned set of SNPs belonging to each component (see above), we found the closest gene to each of these SNPs using the findClosestGene function from the “ACME” R package^10^ (see URLs). This yielded the set of genes belonging to each component.

We calculated the tissue-specific enrichment in each component by dividing the fraction of genes in the component that are in the gene set by the number of genes in the component multiplied by the fraction of all genes in the gene set (i.e. the expected number of genes in the component that are in the gene set). We applied Fisher’s exact test with a Bonferroni correction for the number of tissues.

### Replication r^2^ and variance explained

We define the *replication r^2^* as the squared correlation between true and estimated effect sizes:

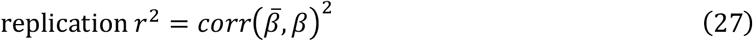

where 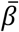 is an estimate of the marginal effect sizes and *β* is the vector of true marginal effect sizes for some trait. The replication r^2^ can be estimated using either a within-sample or an out-of-sample approach. The within-sample replication r^2^ is calculated for a posterior-mean estimator, which should (if unbiased) have the property that

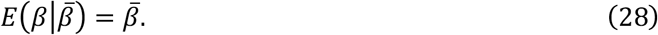

If this property holds, then 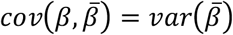 and

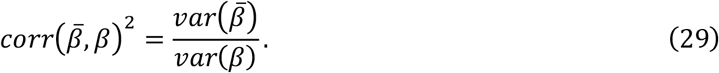

We normalize the summary statistics so that the denominator is 1, and we define the *predicted replication r^2^* as the sample variance of the posterior-mean effect sizes:

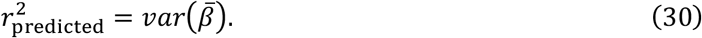

The replication r^2^ can also be quantified out-of-sample. We take summary statistics 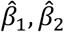 from disjoint cohorts 1 and 2, the training and validation data respectively. We normalize 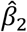 such that 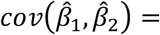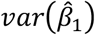, which ensures that the effect sizes are on the same scale. We compute the predicted effect sizes 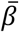 from 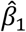. Then, we compute the *observed replication r^2^* as:

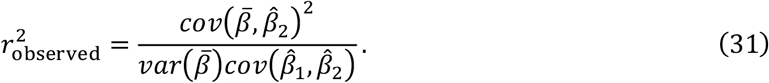

Note that the denominator contains 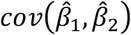 instead of 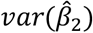 to avoid dilution by noise in the validation data. Using the latter instead would cause all methods to have lower replication r^2^, by a factor that depends on the signal-to-noise ratio in the validation summary statistics.

Replication r^2^ can be defined for any subset of SNPs. However, we choose to report a different metric for subsets of SNPs, the *variance explained*, which is larger than the replication r^2^ when the subset of SNPs is enriched for effect-size variance compared with the rest of the genome. In particular, the variance explained is defined as:

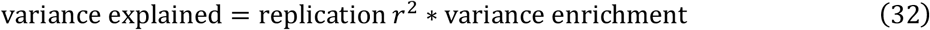

where the variance enrichment of a set of SNPs *A* is:

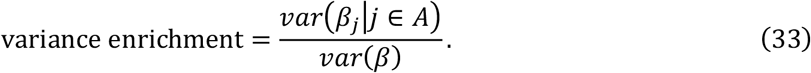

For the set of all SNPs, the variance enrichment is one, and the variance explained is equal to the replication r^2^.

We compare predicted vs. observed variance explained as a way of quantifying bias in the posterior-mean effect sizes for PDR and for MTAG/BLUP. Predicted and observed variance explained are computed in the same way as predicted and observed replication r^2^. If posterior-mean effect sizes are biased specifically for large- or small-effect SNPs, but not on average across all SNPs, then the predicted and observed variance explained will deviate from the line *y* = *x* when they are plotted for subsets of SNPs.

## Simulations

