## Supplementary tables and figures for "Shared components of heritability across genetically correlated traits"

(a) **Asthma Cluster**

| marginal effect size rg | causal effect size rg | Traits |
|---|---|---|
| 0.1497 | 0.1298 | Asthma-BMI |
| 0.6932 | 0.6654 | Asthma-Eczema |
| -0.074 | -0.0806 | BMI-Eczema |
| 0.4464 | 0.4611 | Asthma-Eosinophils |
| 0.107 | 0.1304 | BMI-Eosinophils |
| 0.3172 | 0.3321 | Eczema-Eosinophils |
| -0.3333 | -0.368 | Asthma-FEV1/FVC |
| 0.1527 | 0.1366 | BMI-FEV1/FVC |
| -0.0865 | -0.1098 | Eczema-FEV1/FVC |
| -0.0636 | -0.0851 | Eosinophils-FEV1/FVC |

(b) **Metabolic Cluster**

| marignal effect size rg | causal effect size rg | Traits |
|---|---|---|
| 0.5603 | 0.5633 | T2D-BMI |
| 0.6306 | 0.6878 | T2D-FG |
| 0.2625 | 0.262 | BMI-FG |
| 0.6696 | 0.7459 | T2D-A1C |
| 0.2838 | 0.297 | BMI-A1C |
| 0.5893 | 0.6001 | FG-A1C |
| 0.4645 | 0.492 | T2D-HC |
| 0.2833 | 0.3088 | BMI-HC |
| 0.2256 | 0.2575 | FG-HC |
| 0.3691 | 0.3891 | A1C-HC |
| 0.2746 | 0.2373 | T2D-WHR |
| -0.0754 | -0.0621 | BMI-WHR |
| 0.162 | 0.1623 | FG-WHR |
| 0.1872 | 0.1612 | A1C-WHR |
| 0.2728 | 0.2533 | HC-WHR |

(c) **Coronary Cluster**

| marginal effect size rg | causal effect size rg | Traits |
|---|---|---|
| 0.3302 | 0.4023 | CAD-TG |
| -0.2648 | -0.3178 | CAD-HDL |
| -0.5766 | -0.5622 | TG-HDL |
| 0.5114 | 0.6001 | CAD-HT |
| 0.2391 | 0.2197 | TG-HT |
| -0.2385 | -0.1853 | HDL-HT |
| 0.2772 | 0.3643 | CAD-BMI |
| 0.2769 | 0.2581 | TG-BMI |
| -0.359 | -0.3133 | HDL-BMI |
| 0.3758 | 0.3761 | HT-BMI |
| 0.6243 | 0.6597 | CAD-HC |
| 0.5785 | 0.5214 | TG-HC |
| -0.1598 | -0.1007 | HDL-HC |
| 0.5308 | 0.5405 | HT-HC |
| 0.2963 | 0.3463 | BMI-HC |

**Supplementary Table 1 Marginal and causal effect size correlations.** Comparison of marginal and causal effect size correlations for (a) the asthma cluster, (b) the metabolic cluster, and (c) the coronary cluster. Causal effect-size correlations were computed using LD score regression. Marginal effect-size correlations were computed by subtracting the LD score regression intercept matrix (an estimate of the covariance in the marginal effect-size residuals) from the sample covariance matrix of the summary statistics. The resulting correlations are expected to be similar, except if SNPs with larger or smaller correlations have different amounts of LD. If SNPs with more highly correlated effect sizes had more LD, then marginal effect sizes would be more correlated than causal effect sizes. The estimates do not suggest any such effect.

| | Preferred Model | Factor-like Models | | | | | | | | | |
|---|---|---|---|---|---|---|---|---|---|---|---|
| # cpts | 3 | 1 | 2 | 3 | 4 | 5 | 6 | 7 | 8 | 9 | 10 |
| RSS | 0.4160 | 10.8908 | 4.1382 | 2.3173 | 1.6270 | 1.2545 | 1.2104 | 1.0493 | 0.7931 | 0.8491 | 0.7244 |

**Supplementary Table 2 Objective function values for a preferred model with generic pleiotropic components compared with models with factor-like components.** These results are for models fit to the coronary trait cluster. The preferred model, consisting of three generic pleiotropic components, corresponds to the model used in our main analyses. We compared this model against models with factor-like pleiotropic components with rank-one pattern matrices. The objective function value (RSS, residual sum of squares) for the factor-like models improved as the number of components increased, as expected; however, the RSS for the factor-like models remained larger than that for the preferred model, indicating that full-rank pleiotropic components it the data better than factor-like, rank-one components.

**(a) PDR-derived**

| Column1 | PDR | SEM | SVD |
|---|---|---|---|
| 1 rank-one | 0.0179 | 0.0625 | 0.0867 |
| 2 rank-one | 0.0188 | 0.292 | 0.5689 |
| 3 rank-one | 0.0263 | 0.6405 | 0.8147 |

**(b) Genomic SEM-derived**

| Column1 | PDR | SEM | SVD |
|---|---|---|---|
| 2 rank-one | 0.0469 | 0.0261 | 0.271 |

**(c) SVD-derived**

| Column1 | PDR | SEM | SVD |
|---|---|---|---|
| 2 rank-one | 0.0244 | 0.198 | 0.1009 |

**Supplementary Table 3 Numerical values for method comparison analyses.** Method comparison results are from true models based on (a) PDR (see Figure 1a), (b) genomic SEM (see Supplementary Figure 6a), and (c) SVD (see Supplementary Figure 6b).

| Phenotype | Cluster(s) | Analysis | Data Source | Sample Size |
|---|---|---|---|---|
| **Asthma** | Asthma | Primary analysis | UKB | 458,699 |
| **BMI** | Asthma, T2D, CAD | Primary analysis, Collider bias | UKB | 457,824 |
| **Eczema** | Asthma | Primary analysis | UKB | 458,699 |
| **Eosinophil Count** | Asthma | Primary analysis | UKB | 439,938 |
| **FEV1/FVC** | Asthma | Primary analysis | UKB | 371,949 |
| **T2D** | T2D | Primary analysis | UKB | 459,324 |
| **Fasting Glucose** | T2D | Primary analysis | UKB | 393,767 |
| **HbA1c** | T2D | Primary analysis | UKB | 411,840 |
| **High Cholesterol** | T2D, CAD | Primary analysis | UKB | 459,324 |
| **WHRadjBMI** | T2D | Primary analysis, Collider bias | UKB | 458,417 |
| **CAD** | CAD | Primary analysis | CARDIoGRAM | 77,210 |
| **Triglycerides** | CAD | Primary analysis | UKB | 431,952 |
| **HDL** | CAD | Primary analysis | UKB | 397,612 |
| **Hypertension** | CAD | Primary analysis | UKB | 458,554 |
| **Asthma** | Asthma | Out-of-sample replication | TAGC | 127,669 |
| **T2D** | T2D | Out-of-sample replication | DIAGRAM | 60,786 |
| **WHRunadj** | None | Collider bias | GIANT | 138,387 |
| **Years of Education** | None | Adding EA to coronary and asthma clusters | UKB | 454,813 |

**Supplementary Table 4 Summary statistics used in all analyses.** Primary analysis data was used in the main analyses to get the heat maps, scatter plots, and expected replication $r^2$ and variance explained, as well as to get tissue-specific enrichments. Out-of-sample replication refers to the analyses performed to obtain observed replication $r^2$. This table includes the phenotype, the trait cluster(s) to which that phenotype belongs, the analysis or analyses that used the data, the data source, and the sample size. See URLs to access the summary statistics.

| Trait cluster | 1 vs. 2 | 2 vs. 3 | 3 vs. 4 |
|---|---|---|---|
| Coronary | 3.68E-50 | 8.07E-05 | 9.99E-01 |
| Asthma | 2.28E-30 | 4.70E-22 | 1.84E-02 |
| Metabolic | 2.22E-47 | 7.52E-07 | 9.99E-01 |

**Supplementary Table 5 P-values comparing models with different numbers of pleiotropic components**. Significant p-values indicate that the larger model fits better. These p-values were calculated using a tolerance parameter of 0.005 (see “Regression weights” in Methods).

(a)

| Coronary cluster | p-values |
|---|---|
| **CAD** | 4.38E-01 |
| **TG** | 8.06E-01 |
| **HDL** | 2.42E-01 |
| **Hypertension** | 4.38E-01 |
| **BMI** | 6.43E-01 |
| **High Cholesterol** | 9.95E-01 |

(b)

| Asthma cluster | p-values |
|---|---|
| **Asthma** | 7.13E-01 |
| **BMI** | 8.63E-01 |
| **Eczema** | 9.15E-03 |
| **Eosinophil count** | 4.44E-04 |
| **FEV1/FVC** | 2.47E-01 |

(c)

| Metabolic cluster | p-values |
|---|---|
| **T2D** | 4.90E-01 |
| **BMI** | 6.47E-01 |
| **Glucose** | 9.04E-01 |
| **HbA1c** | 9.40E-16 |
| **High Cholesterol** | 3.11E-02 |
| **WHRadjBMI** | 1.00E+00 |

**Supplementary Table 6 P-values for trait-specific components for preferred models in each cluster.** Each preferred model consisted of trait-specific components for each trait and three generic pleiotropic components. The null models were created by removing one trait-specific component at a time and the alternative was the preferred model. Results are shown for the (a) asthma cluster, (b) metabolic cluster, and (c) coronary cluster. These p-values were calculated using a tolerance parameter of 0.005 (see "Regression weights" in Methods).

| Cpt | tissue | enrichment pval | enrichment magnitude | # expected genes | # observed genes |
|---|---|---|---|---|---|
| BMI | Brain Caudate (basal ganglia) | 4.84E-07 | 1.14 | 195 | 222 |
| BMI | Brain Frontal Cortex (BA9) | 1.72E-06 | 1.13 | 200 | 226 |
| BMI | Brain Amygdala | 1.97E-06 | 1.13 | 199 | 225 |
| BMI | Brain Anterior cingulate cortex (BA24) | 2.24E-06 | 1.13 | 198 | 224 |
| BMI | Brain Putamen (basal ganglia) | 3.31E-06 | 1.13 | 196 | 221 |
| BMI | Brain Hypothalamus | 5.17E-06 | 1.12 | 199 | 224 |
| BMI | Brain Hippocampus | 1.29E-05 | 1.12 | 199 | 223 |
| BMI | Brain Cortex | 2.09E-05 | 1.12 | 196 | 219 |
| BMI | Brain Nucleus accumbens (basal ganglia) | 3.00E-05 | 1.12 | 193 | 216 |
| BMI | Brain Substantia nigra | 8.73E-05 | 1.11 | 203 | 225 |
| BMI | Brain Spinal cord (cervical c-1) | 5.65E-04 | 1.09 | 200 | 219 |
| Cholesterol | Adipose Visceral (Omentum) | 7.88E-04 | 1.65 | 24 | 40 |

**Supplementary Table 7 Significant tissue-specific enrichments for the CAD cluster.** Significance was determined based on a Bonferroni threshold of 0.05 corrected for the number of tissues, 53. See Supplementary Table 14 for all tissue enrichments and p-values.

| cpt | tissue | enrichment pval | enrichment magnitude | # expected genes | # observed genes |
|---|---|---|---|---|---|
| Pulmonary | Esophagus Gastroesophageal Junction | 1.96E-06 | 1.74 | 36 | 64 |
| Pulmonary | Artery Tibial | 8.72E-06 | 1.66 | 39 | 66 |
| Pulmonary | Esophagus Muscularis | 3.51E-05 | 1.64 | 35 | 59 |
| BMI | Brain Hippocampus | 3.02E-04 | 1.09 | 191 | 209 |
| BMI | Brain Amygdala | 4.38E-04 | 1.09 | 188 | 205 |
| Inflammatory | Spleen | 7.41E-08 | 2.21 | 20 | 46 |
| Inflammatory | Small Intestine Terminal Ileum | 8.05E-04 | 1.65 | 24 | 40 |

**Supplementary Table 8 Significant tissue-specific enrichments for the asthma cluster.** Significance was determined based on a Bonferroni threshold of 0.05 corrected for the number of tissues (53). See Supplementary Table 14 for all tissue enrichments and p-values.

| cpt | tissue | enrichment pval | enrichment magnitude | # expected genes | # observed genes |
|---|---|---|---|---|---|
| BMI | Brain Hypothalamus | 1.88E-04 | 1.11 | 194 | 216 |
| BMI | Brain Caudate (basal ganglia) | 2.06E-04 | 1.11 | 189 | 210 |
| BMI | Brain Nucleus accumbens (basal ganglia) | 2.95E-04 | 1.11 | 191 | 212 |
| BMI | Brain Frontal Cortex (BA9) | 3.30E-04 | 1.11 | 190 | 211 |
| BMI | Brain Putamen (basal ganglia) | 3.72E-04 | 1.10 | 194 | 215 |
| BMI | Brain Amygdala | 5.15E-04 | 1.10 | 192 | 212 |
| BMI | Brain Anterior cingulate cortex (BA24) | 7.88E-04 | 1.10 | 189 | 208 |

**Supplementary Table 9 Significant tissue-specific enrichments for the metabolic cluster.** Significance was determined based on a Bonferroni threshold of 0.05 corrected for the number of tissues, 53. See Supplementary Table 14 for all tissue enrichments and p-values.

**Supplementary Table 10 PDR model parameters for coronary, asthma, and metabolic clusters.** Data for the PDR models with three pleiotropic components fit to the coronary, asthma, and metabolic clusters, including trait names, scalars, factor weights (theta), mixture weights (ww), covariance matrix (cov), and pattern matrix. See Supplementary_Tables.xlsx S10a-c for the parameters from the models fit to the (a) coronary, (b) asthma, and (c) metabolic clusters.

**Supplementary Table 11 Numerical values for models fit to the coronary and asthma clusters with EA added as an additional trait.** See Supplementary_Tables.xlsx S11a-b for the numerical values comprising the upper triangular elements of each component covariance and correlation matrix for the (a) coronary and (b) asthma trait clusters.

**Supplementary Table 12 Pruned SNP information for coronary, asthma, and metabolic clusters.** Data for the ~11K SNPs that remained after LD pruning, including the SNP rsID, posterior mean effect sizes on each trait in the cluster, posterior scalars for each component, and the component to which that SNP was assigned. The posterior mean effect sizes for each trait and posterior scalars for each component appear in the same order as described in Supplementary Table 10. See Supplementary_Tables.xlsx S12a-c for the data from the (a) coronary, (b), asthma, and (c) metabolic clusters.

**Supplementary Table 13 Numerical values for replication $r^2$ and per-SNP variance explained analyses.** The numerical values of replication $r^2$ and variance explained plotted in Figure 4 are presented in Supplementary_Tables.xlsx S13a-c. Note that the per-SNP variance explained values are on the log10 scale. Expected and observed replication $r^2$ and variance explained are shown for the (a) asthma and (b) metabolic clusters. (c) Only the expected replication $r^2$ is shown for the coronary cluster, since out-of-sample replication data for CAD were not available.

**Supplementary Table 14 All tissue enrichments and p-values for the coronary, asthma, and metabolic clusters.** See Supplementary_Tables.xlsx S14a-c for the enrichment results and p-values for all 53 tissues in the (a) coronary, (b) asthma, and (c) metabolic clusters. Values for each component are sorted by enrichment p-value in ascending order. Note that the majority of the enrichments are nonsignificant. An enrichment p-value of 1 corresponds to the case when there are zero genes observed in the component that belong to that gene set, including when the expected number of genes is also zero.

**Supplementary Table 15 Parameters of models used in method comparison analysis.** See Supplementary_Tables.xlsx S15a-e for the traits, scalars, factor weights (theta), mixture weights (ww), covariance matrix (cov), and pattern matrix used in (a-c) each of the 1-3 “factor-like” rank-one component models inferred from PDR, (d) the 2 component model designed to favor genomic SEM and (e) the 2 component model designed to favor SVD.

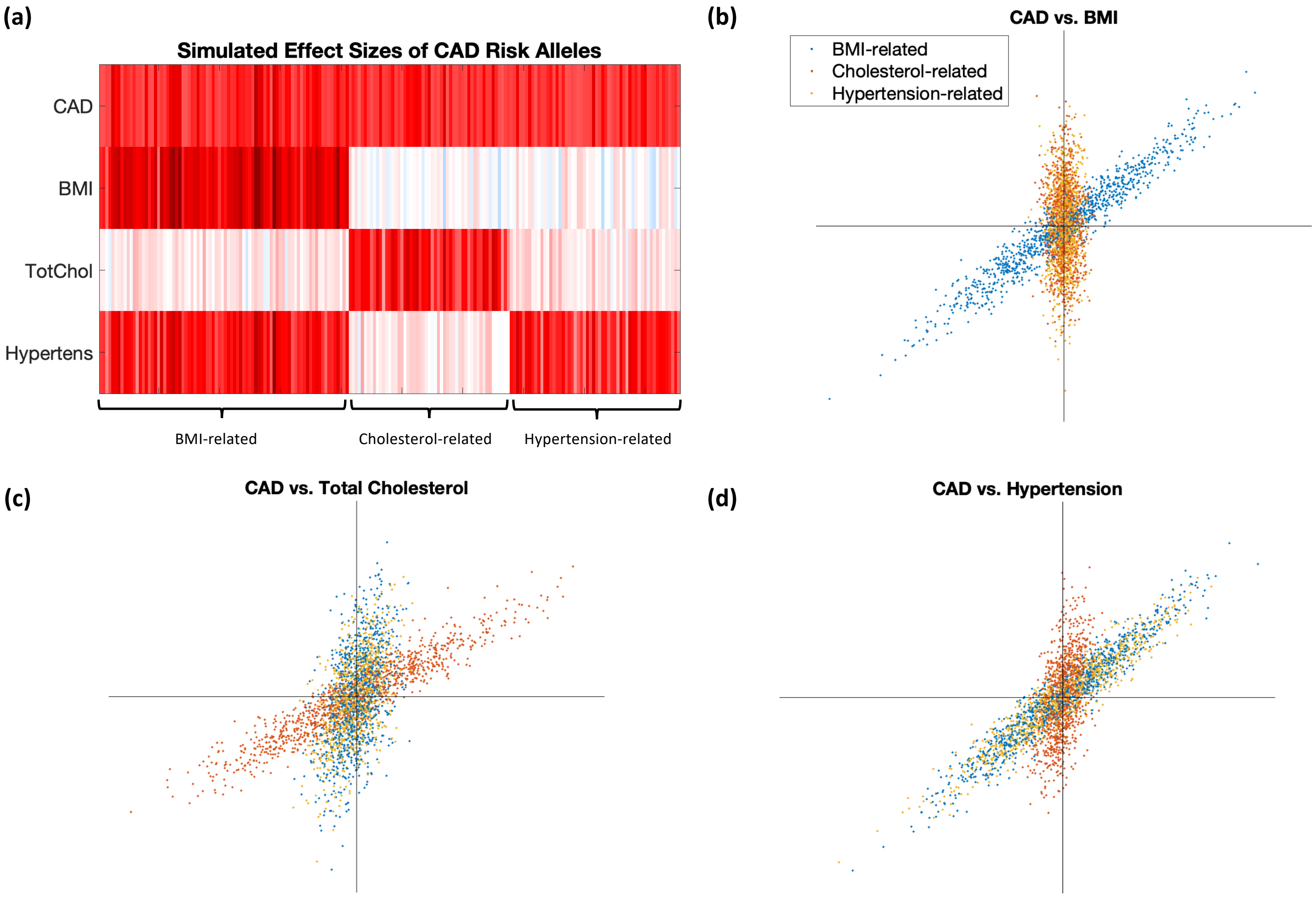


**Supplementary Figure 1 Illustration of how PDR links SNPs, components, and traits using simulated data.** (a) Heatmap showing the simulated effect sizes of each the SNPs on each trait. These SNPs form three clusters, BMI-related, Cholesterol-related, and Hypertension-related, in which they demonstrate similar patterns of association. Each of these clusters are distinguished by the set of traits they primarily affect. (b-d) Scatter plots of the SNP effect sizes on three trait pairs (titles are shown as y-axis trait vs. x-axis trait). Each of the scatterplots reflects the simulated effect sizes shown in the heatmap. (b) SNPs in the BMI-related cluster affect both CAD and BMI with positively correlated effect sizes, whereas those in the cholesterol-related and hypertension-related clusters affect CAD but not BMI. (c) SNPs in the cholesterol-related component affect both CAD and total cholesterol with positively correlated effects, but SNPs in the BMI-related and hypertension-related components affect CAD but not total cholesterol. (d) SNPs in the BMI-related and hypertension-related components affect both CAD and hypertension with positively correlated effects, and cholesterol-related SNPs affect CAD but not hypertension. The total effect size distribution is the sum of the SNP effects within each component. Since the SNPS here are simulated to belong to a single component, the total effect size distribution equivalent to the superposition of the three colored scatterplots.

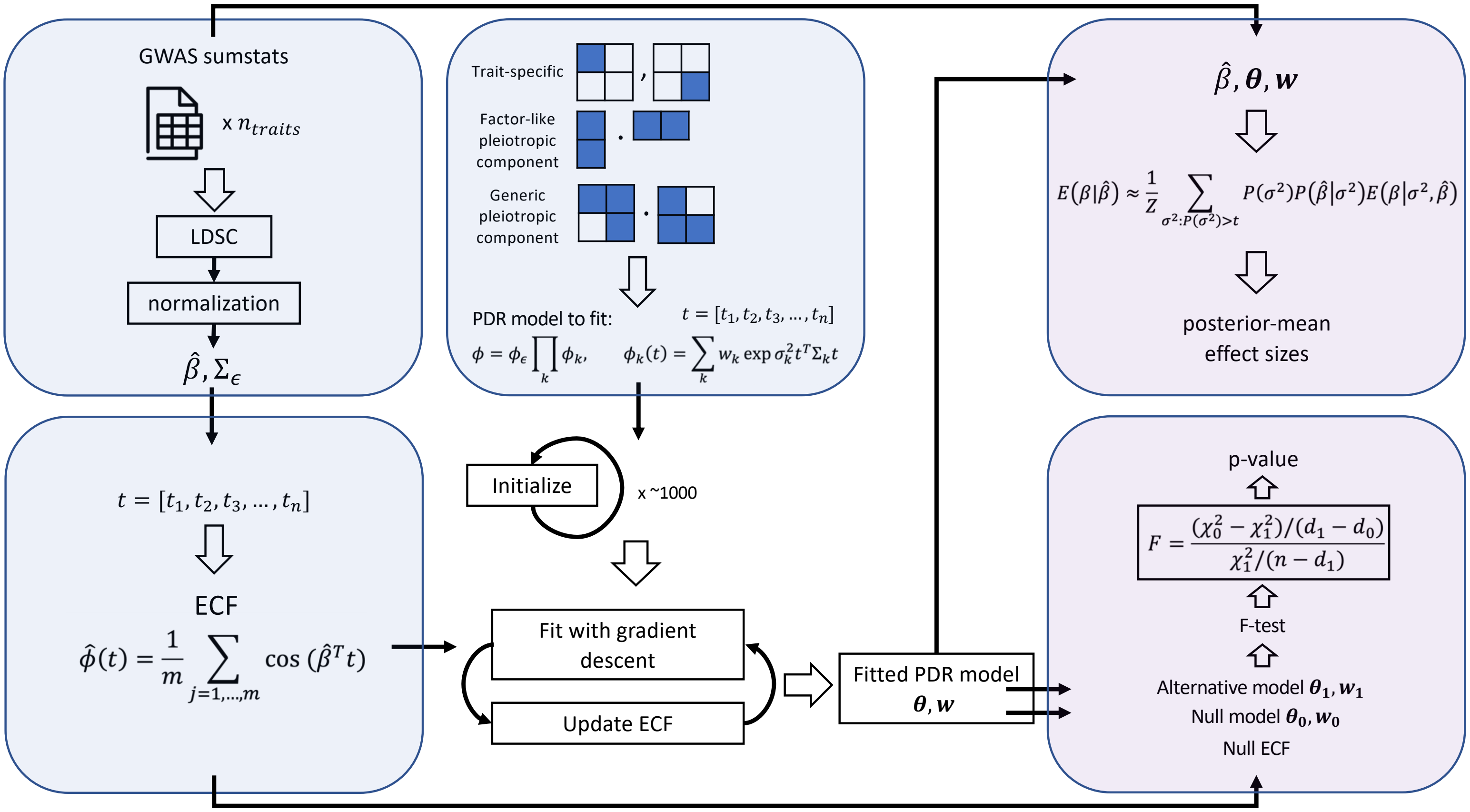


**Supplementary Figure 2 Overview of PDR.** The blue boxes correspond to the major elements used in PDR: the data, the model, and the empirical characteristic function (ECF). The GWAS summary statistics are loaded and preprocessed using LDSC and normalization, and they are used to calculate the ECF. A model scaffold is built using a user-specified combination of trait-specific, factor-like pleiotropic, and generic pleiotropic components; other component types are also possible. Then the model is fit starting with several random initializations, performing just three gradient descent steps for each initialization, and the one with the best objective function value is chosen. PDR iteratively fits this model using gradient descent and updates the ECF with a new regression weights matrix until convergence. The fitted model is used in downstream analyses. These include calculating posterior-mean effect sizes using the data and the fitted model as inputs, and hypothesis testing to compare how well a null and an alternative model fit the data using an F-test.

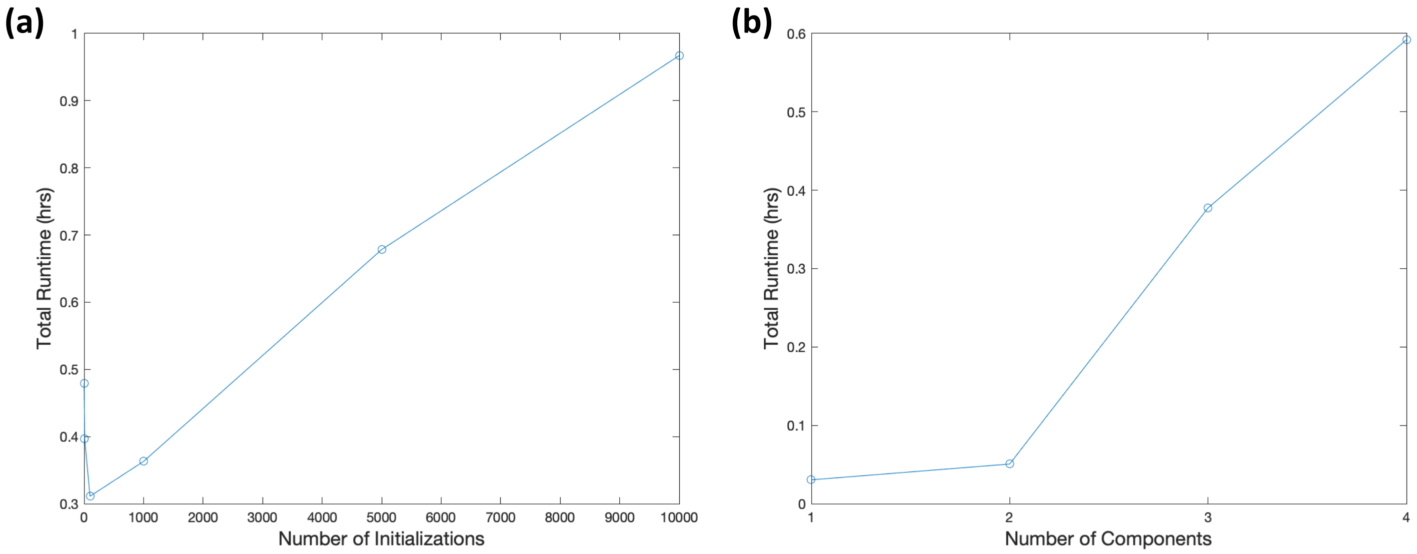


**Supplementary Figure 3 PDR runtime measured for varied number of initializations and components.** Total runtime is measured as the sum of the total time it takes to complete the initialization step and the total time it takes to complete the iterative process of fitting the model with gradient descent and updating the ECF. Each point represents the average runtime of five replicates. (a) Runtime as a function of number of initializations. This was for a model with six traits and three components. Note that the runtime decreased from 1 to 100 initializations. This was because the runtime for the gradient descent step was much larger for these settings despite having fewer initializations, likely because there were too few initializations to get sufficiently close to the optimum, necessitating more gradient descent steps to converge on a solution. (b) Runtime as a function of number of components. The number of traits was fixed at six, and the number of initializations was held at 100.

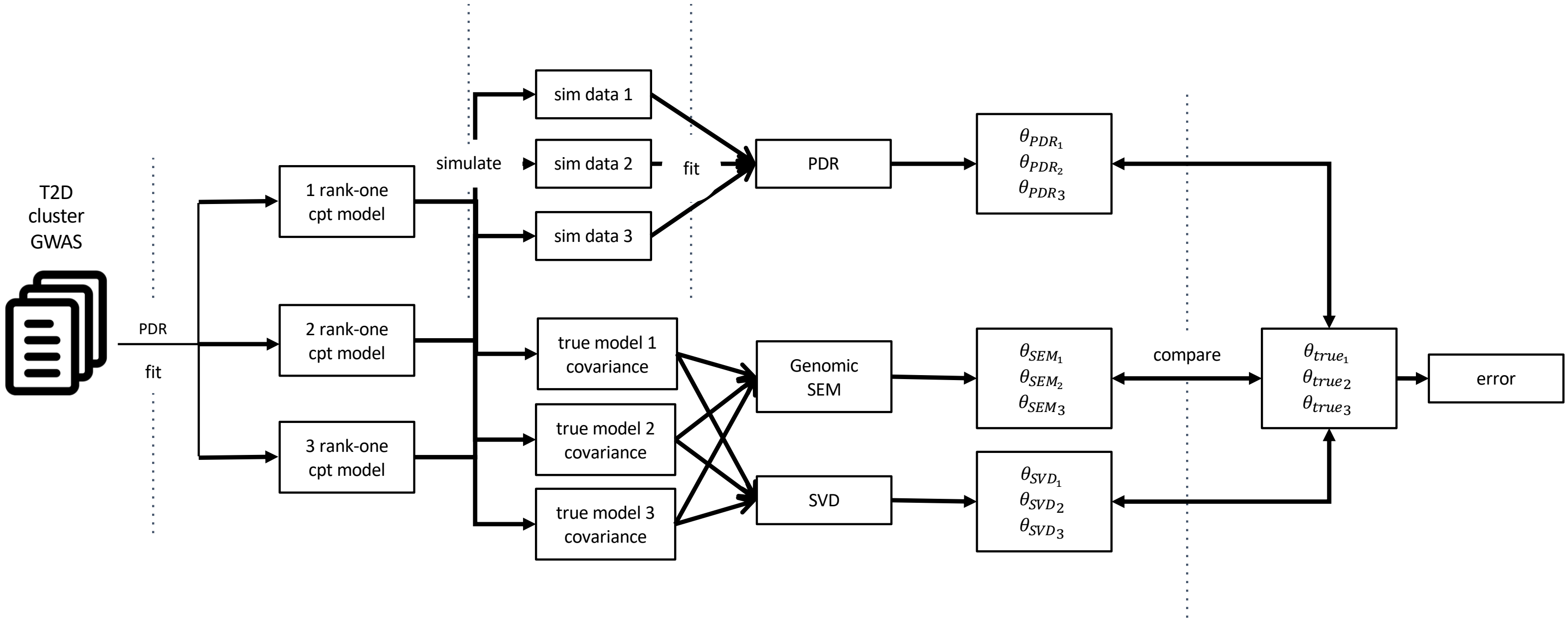


**Supplementary Figure 4 Schematic for method comparison simulations.** Three PDR models consisting of one, two, and three rank-one components were fit to the cluster of metabolic traits. Although we found that "partially correlated" components with full-rank pattern matrices fit the data better in general (Supplementary Table 2), we used factor-like components for the purpose of model comparisons because they match the structure that genomic SEM and SVD aim to infer. These components can be interpreted as latent biological variables affecting each trait with effect sizes proportional to the factor weights. The true model covariance matrices were used to calculate the factor weights, $\theta$, for SEM and SVD, whereas the factor weights for PDR were calculated by fitting PDR models to data simulated from the three true models.

| | Component 1 | | | | Component 2 | | | | Component 3 | | | |
|---|---|---|---|---|---|---|---|---|---|---|---|---|
| Trait | True factor weights | PDR | SEM | SVD | True factor weights | PDR | SEM | SVD | True factor weights | PDR | SEM | SVD |
| T2D | 0.017 | 0.014 | 0.118 | 0.425 | 0.567 | 0.572 | 0.567 | 0.203 | 0.589 | 0.584 | 0.153 | 0.567 |
| BMI | -0.369 | -0.360 | 0 | -0.053 | -0.134 | -0.138 | 0.085 | -0.400 | 0.566 | 0.571 | 0.910 | -0.134 |
| FG | 0.012 | 0.004 | 0 | 0.240 | 0.337 | 0.352 | 0.509 | 0.178 | 0.298 | 0.315 | 0 | 0.337 |
| A1C | -0.049 | -0.053 | -0.115 | 0.263 | 0.451 | 0.446 | 0.635 | 0.389 | 0.310 | 0.310 | 0 | 0.451 |
| HiChol | 0.557 | 0.558 | 0.778 | 0.533 | 0.258 | 0.243 | 0 | -0.749 | 0.374 | 0.358 | 0.167 | 0.258 |
| WHRadjBMI | 0.743 | 0.746 | 0.606 | 0.637 | 0.526 | 0.521 | 0.099 | 0.232 | 0.093 | 0.093 | -0.348 | 0.526 |

**Supplementary Figure 5 Method comparison for true model with 3 rank-one components.** Comparison of the true model thetas with the optimally matched thetas estimated by PDR, SEM, and SVD, where the true model has three rank-one components. The PDR factor weights were taken from one of the ten replicates, but the error (0.0199) was similar to the mean error across all replicates (mean=0.0263, std=0.0079).

(a)

(b)

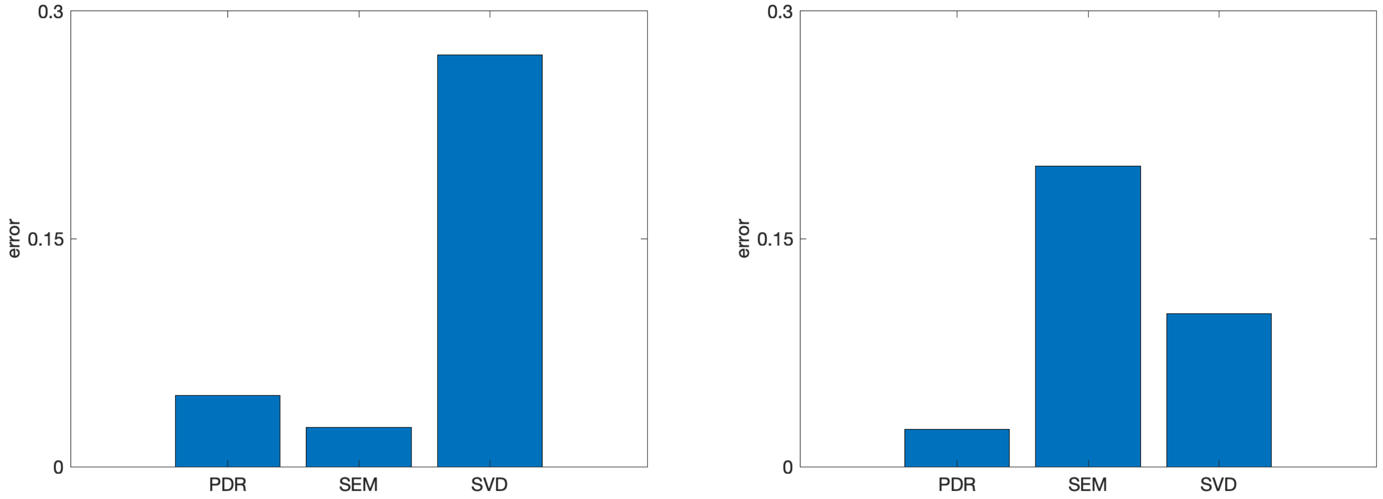


**Supplementary Figure 6 Error for factor weights estimated by PDR, SEM, and SVD on data simulated from true models based on SEM and SVD.** (a) Error for a model based on genomic SEM. (b) Error for a model based on SVD. See Supplementary Table 3b-c for numerical results.

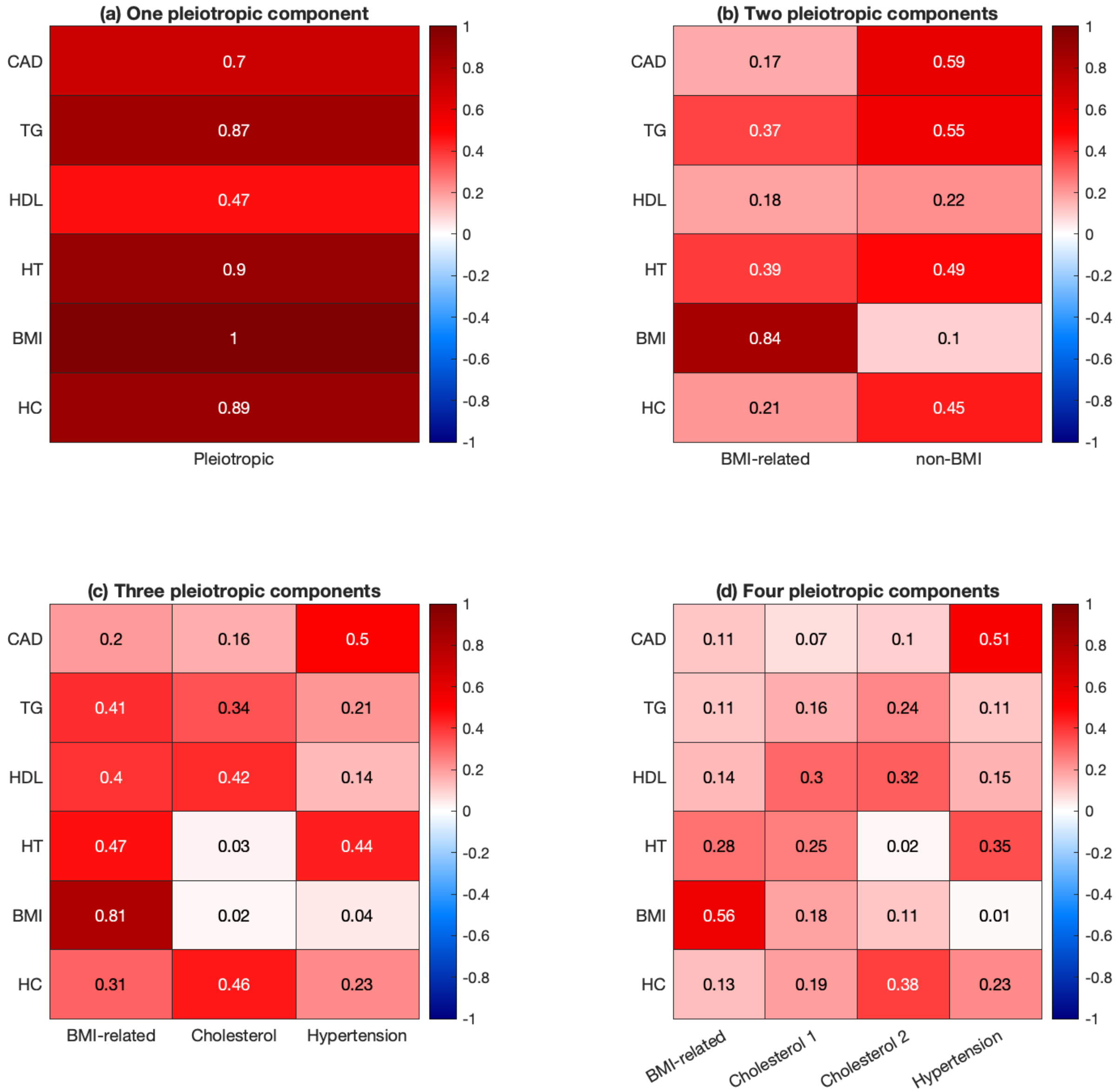


**Supplementary Figure 7 Variance explained by each component on each trait for the coronary cluster.** These results are for models with 1-4 pleiotropic components.

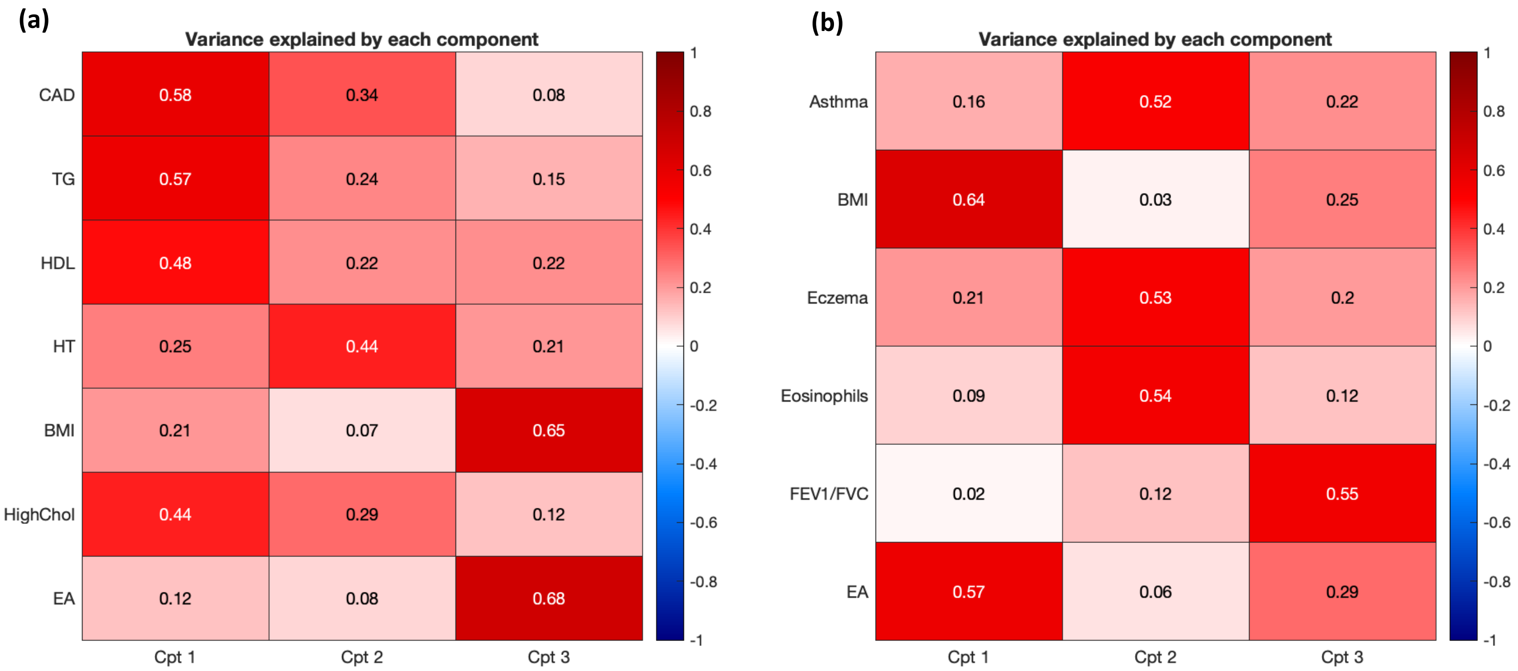


**Supplementary Figure 8 Variance explained for coronary and asthma clusters with educational attainment included as an additional trait.** (a) Coronary cluster with EA demonstrated components with similar trait association patterns as when EA was not included, with the BMI-related component now explaining a large proportion of both BMI and EA variance. The addition of EA did not cause large changes in the correlations between BMI and CAD. The BMI-related component (Cpt 3 above) had a CAD-BMI correlation of 0.92 and a CAD-EA correlation of -0.58 (see Supplementary Table 11 for all component covariance and correlation matrices), demonstrating that adding EA did not decrease the correlation between BMI and CAD, which might have been expected if BMI was merely a proxy for the effect of EA. (b) We also did a similar analysis for the asthma cluster and observed similar results. After EA was included, the former BMI-related component (Cpt 1 above) explained a large proportion of both BMI and EA variance. It had an asthma-BMI correlation of 0.51 and an asthma-EA correlation of -0.20, demonstrating that adding EA did not cause a drastic reduction in the correlation between asthma and BMI.

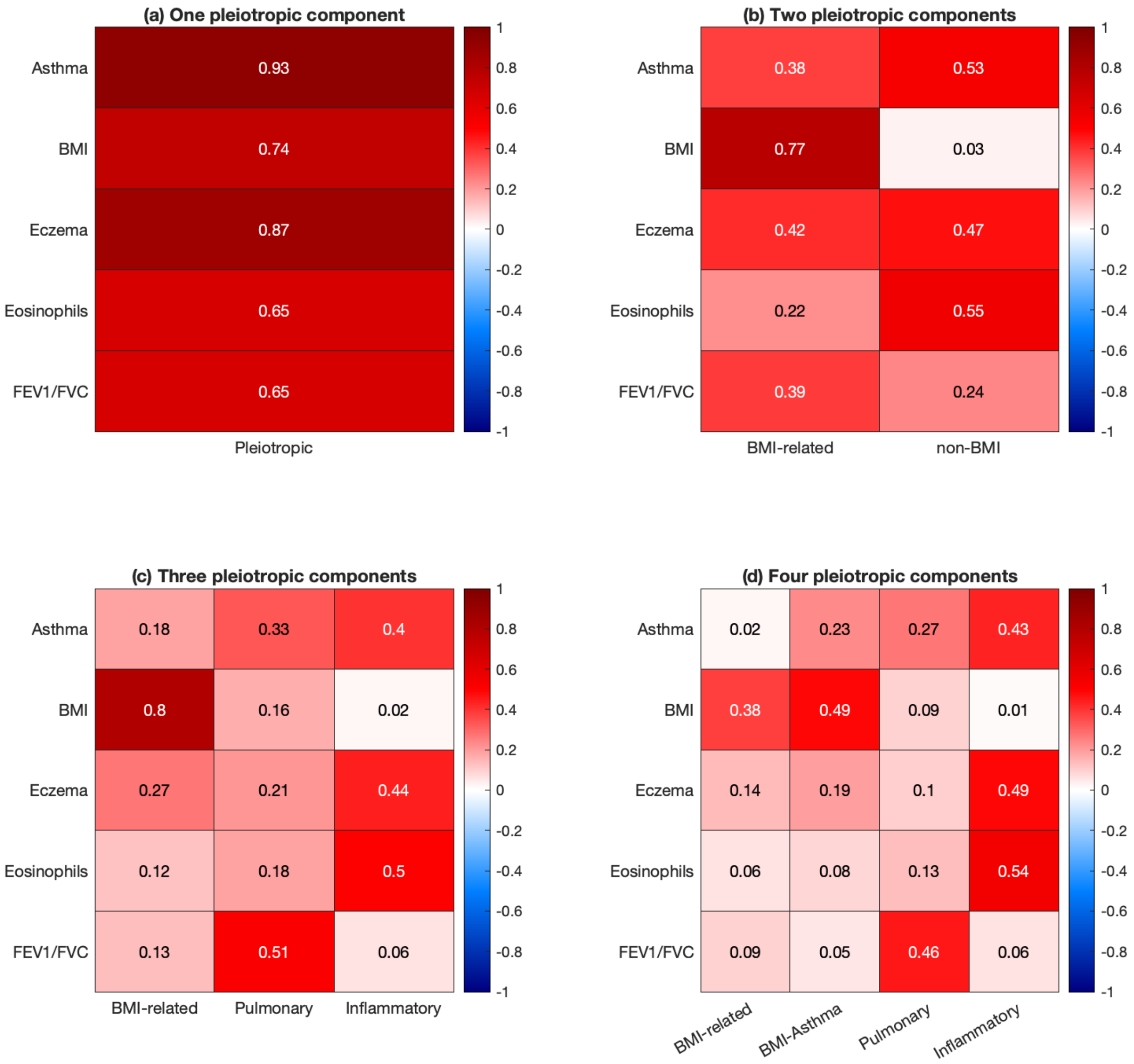


**Supplementary Figure 9 Variance explained by each component on each trait for the asthma cluster.** These results are for models with 1-4 pleiotropic components.

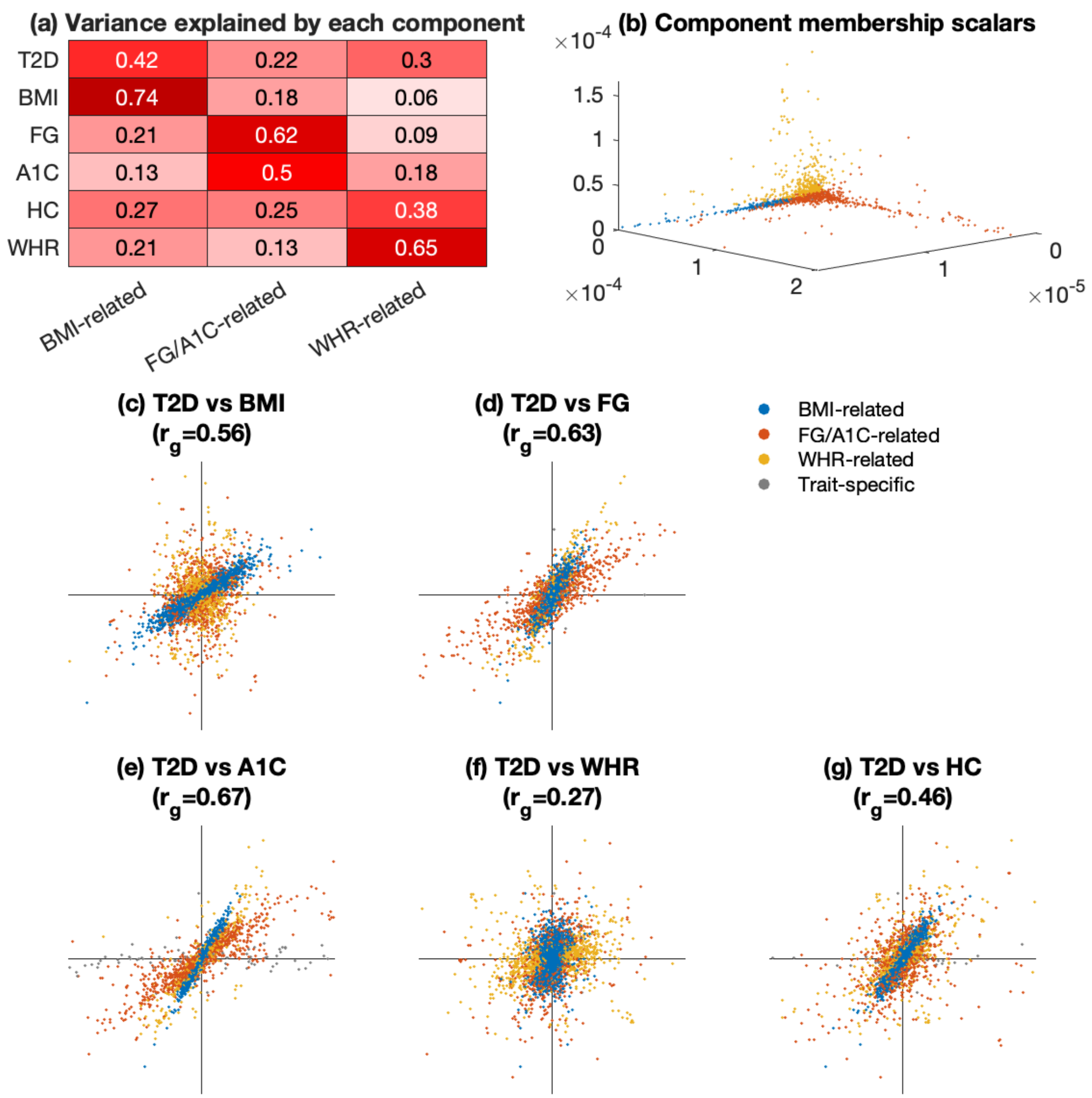


**Supplementary Figure 10 Shared heritability components for T2D.** We applied PDR to summary statistics for T2D and five genetically correlated metabolic traits (Supplementary Table 4 and URLs). We performed LD pruning and computed posterior-mean effect sizes of the remaining SNPs (M=11k). (a) Variance explained by the three pleiotropic components for each trait. (b) Posterior-mean component membership scalars for each SNP, colored by for visualization purposes. (c) Posterior-mean effect sizes on T2D (y-axis) and BMI (x-axis); (d) posterior-mean effect sizes on T2D and fasting glucose; (e) posterior-mean effect sizes on T2D and HbA1c; (f) posterior-mean effect sizes on T2D and WHRadj; (g) posterior-mean effect sizes on T2D and high total cholesterol. See Supplementary Tables 10 and 12 for the fitted model parameters and pruned SNP information, respectively. Like the other trait clusters, a model with three pleiotropic components fit better than a model with two or one, and equally well as a model with four (Supplementary Table 5, Supplementary Figure 11). We identified a BMI-related component, an FG- and A1C-related component, and a WHR-related component. HbA1c had a significant trait-specific component (Supplementary Table 6). The BMI pleiotropic component explained the majority of BMI heritability, and it had strongly correlated effects on the other traits except for WHRadj. The FG-A1C component had strongly correlated effects on T2D, fasting glucose and HbA1c; it also had weakly correlated effects on BMI, HC and WHRadj. The WHR-related component most strongly affected WHRadj, high cholesterol and T2D, with weakly correlated effects. We also performed a gene-set enrichment analysis and found that, similar to the CAD and asthma trait clusters, the BMI-related component was significantly enriched in brain tissues; the other components did not have any significant enrichments (Supplementary Table 9).

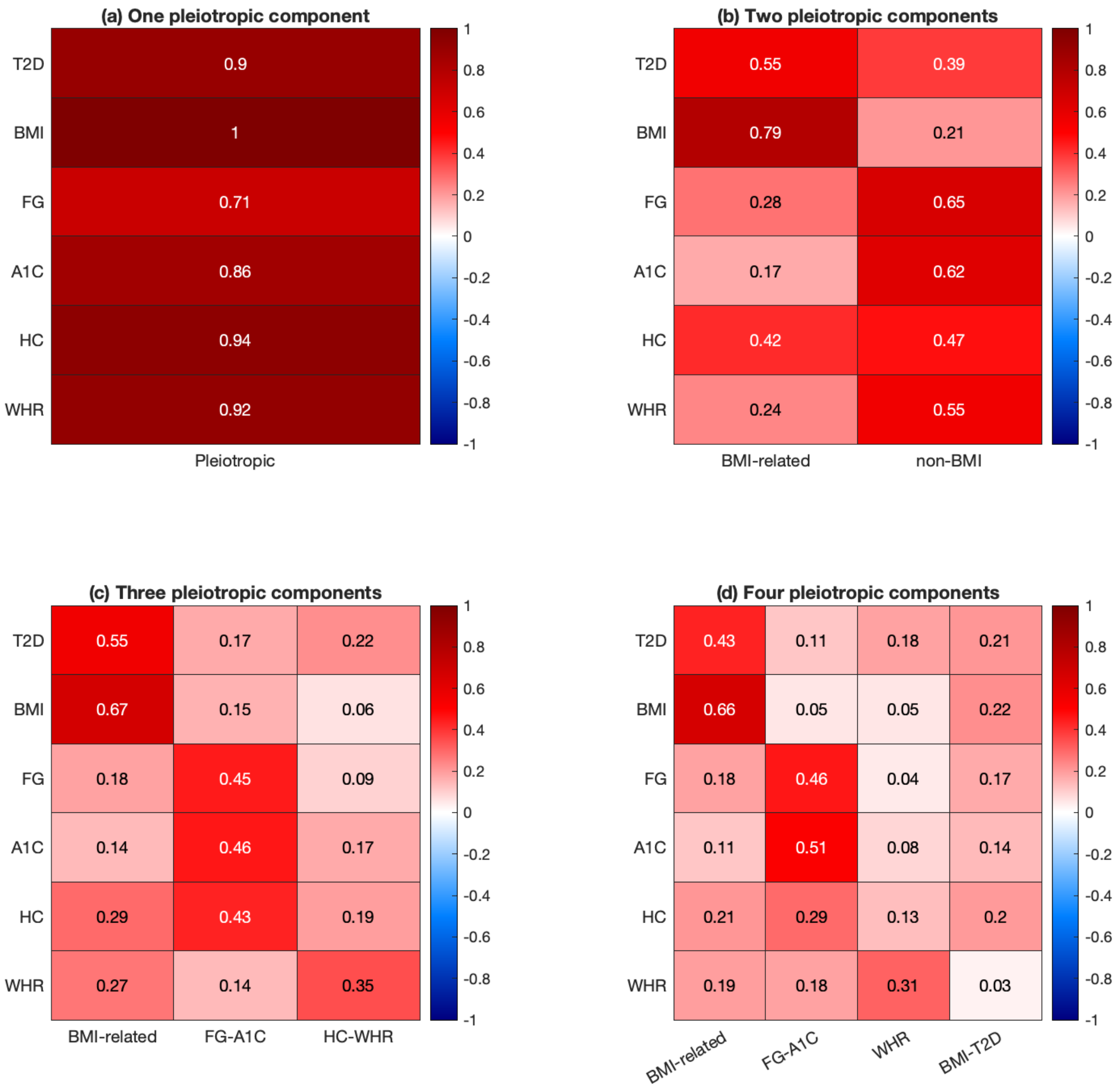


**Supplementary Figure 11 Variance explained by each component on each trait for the metabolic cluster.** These results are for models with 1-4 pleiotropic components.

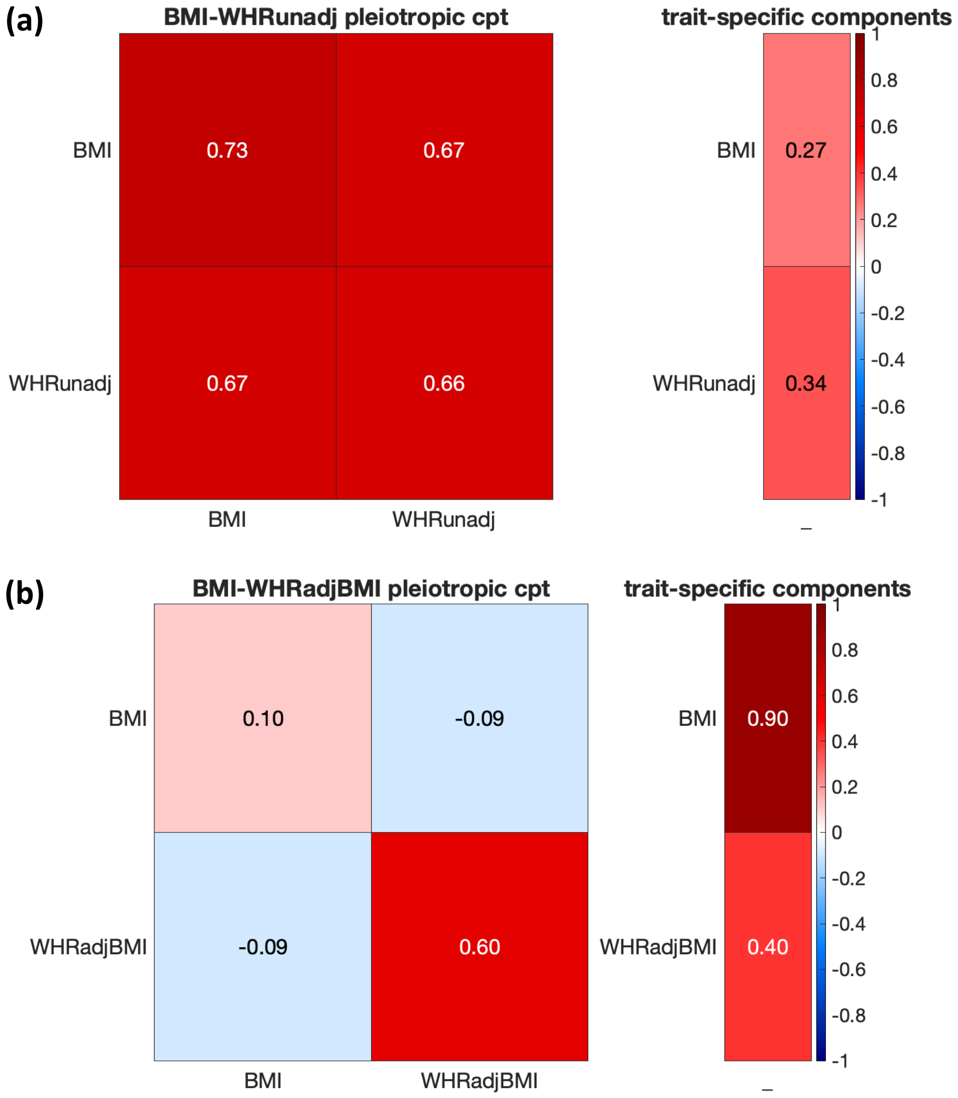


**Supplementary Figure 12 Analyses of BMI and waist-hip ratio to assess collider bias.** We were concerned about performing analyses with waist-hip ratio adjusted for BMI because of potential collider bias: if any SNPs affected BMI but not WHR, then after adjustment for BMI, these SNPs might be spuriously associated with WHRadjBMI. We applied PDR to BMI and waist-hip ratio (WHR) before or after it was adjusted (panels a-b respectively). A model fit to BMI and unadjusted WHR resulted in a component explaining a similar fraction of heritability for both BMI and WHR with nearly perfectly correlated effects. It suggests that there is a large proportion of SNPs that affect both BMI and WHR, which would be corrected by adjusting WHR for BMI. Potentially, the BMI-specific heritability (comprising 27% of variance) might lead to spurious associations with WHRadjBMI. However, fitting a model to BMI and WHRadjBMI yielded a pleiotropic component that was nearly WHRadjBMI-specific, indicating that adjusting WHR for BMI successfully removed the dependency between WHR and BMI without creating substantial spurious associations. Therefore, collider bias is not expected to have a large impact on our results, and we are able to include WHRadjBMI along with BMI in our metabolic cluster analyses.

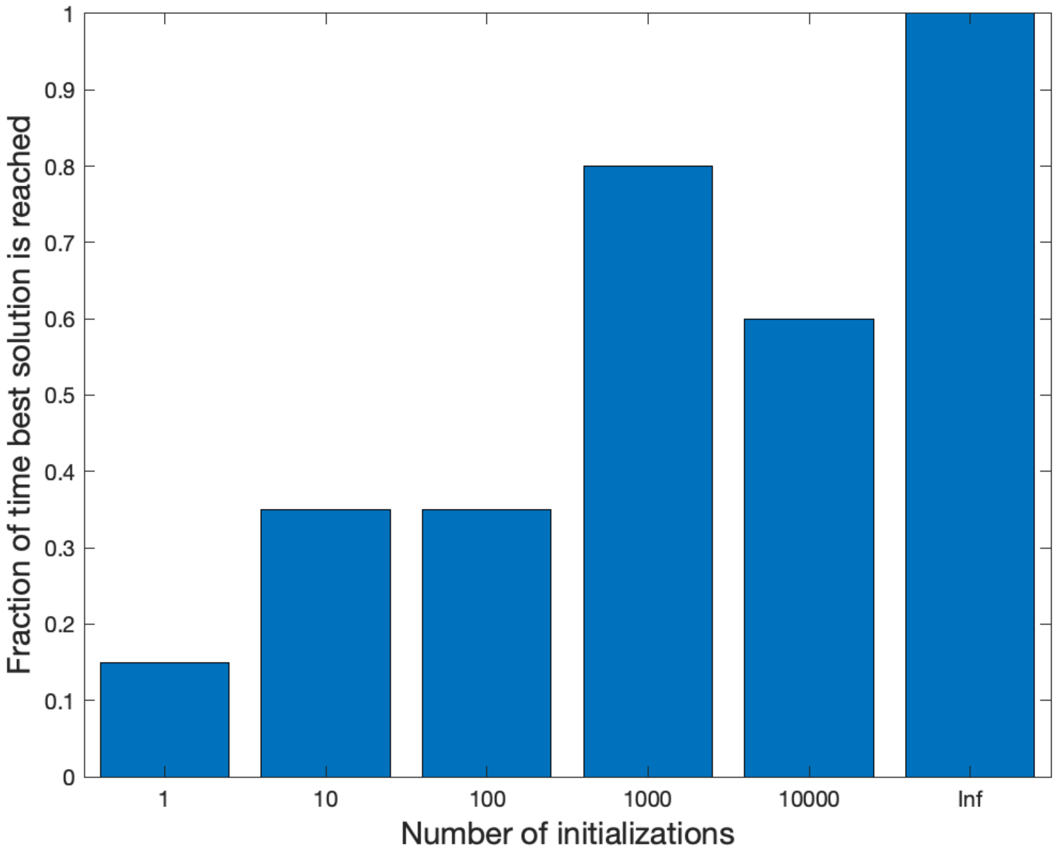


**Supplementary Figure 13 Fraction of best solutions for different numbers of random initializations.** For 1 to 10,000 initializations, parameters were initialized using the 'init_cov' method, which draws values from a multivariate normal distribution with covariance matrix equal to the data covariance. The results for "Inf" initializations were obtained using the 'init_rand' method, which includes a single initialization at the true parameters and several random initializations. This approach approximates the limit of infinite initializations, the set of which would include the true parameters. Twenty replicates were performed for 1, 10, and 100 initializations, and 5 replicates for 1,000, 10,000, and Inf initializations. At 1,000 and 10,000 initializations, we performed fewer replicates due to computational cost.
